# HomoeoGWAS enables experiment-wide homoeolog-group interaction mapping in allopolyploid crops

**DOI:** 10.64898/2026.11.06.764877

**Authors:** Cheng Si, Yujiang Wu, Zhu Sun, Cai Yao, Wei Li, Haiwang Zhang, Lihui Wang, Xuemei Sun, Qiwen Zhong, Shipeng Yang

## Abstract

Allopolyploid crops carry homoeologous gene copies from divergent subgenomes, an evolutionarily defined set of candidate interactions that association studies do not test directly. HomoeoGWAS uses these copies to define the unit of interaction testing, the homoeolog group, and analyses all eligible groups as one family with experiment-wide error calibration, combining three complementary genotype encodings per edge with a structure-aware parametric bootstrap. Family-wise error was controlled in 80-group tests in wheat, cotton and rapeseed under several null models and in full-family tests of all four crops under a structure-aligned global null (379– 17,404 groups). The omnibus outperformed a SNP-by-SNP scan when interaction spanned several genotype dimensions. HomoeoGWAS detected homoeolog-group interactions for heading date in wheat, flowering time in rapeseed and fibre length in cotton; five groups at four loci retained support after regional conditioning. Rapeseed homoeolog coupling showed a weak but reproducible association with homoeologous-exchange frequency. Genome-wide interaction analyses took minutes.

## Introduction

Several major crops, including bread wheat, cotton, oilseed rape and peanut^1–4^, are allopolyploids that combine two or three divergent subgenomes in one nucleus, so that many genes are present as homoeologous copies^5–7^. These copies are not independent: their expression is coordinated and frequently biased towards one subgenome^8–10^, their dosage balance can shape phenotype^11^, and in some species, such as *Brassica napus*, recurrent homoeologous exchanges reshuffle copies between subgenomes^3,12–15^. Allopolyploid genomes therefore carry a biologically defined set of candidate genetic interactions among homoeologous copies.

Standard association workflows do not test this set directly. Genome-wide association studies test one marker or one gene at a time, and the mixed models used in polyploid crops either treat the genome as a single kinship^16–18^ or model allele dosage within a marker^19^. Interaction scans in their general form test all marker pairs^20,21^, a search space that reached 1.96 × 10¹¹ pairs for the 627,255 SNPs mapped to the 17,404 rapeseed homoeolog groups studied here, with a multiplicity and computational cost that make it impractical as a routine analysis^22,23^; recent marginal tests instead aggregate the interactions of each variant with the rest of the genome^24,25^. Subgenome-partitioned variance components with additive-by-additive terms improve genomic prediction in wheat^26,27^, but a variance component estimates an aggregate contribution; it does not test which homoeologous genes interact. What is missing is a calibrated, genome-wide association test whose unit is the homoeolog group itself.

Such a test must meet three requirements. It must be sensitive to interactions of different form, because homoeologs can act through dosage, through shared haplotype structure or through more complex genotype combinations, and a test built for one form can be blind to another. It must control the experiment-wide error rate across thousands to tens of thousands of groups in panels with strong population structure, in which residual variance itself often follows that structure. And it must be fast and reproducible enough to be applied routinely to whole genomes.

Here we present HomoeoGWAS, which uses the evolutionary relationship between homoeologous copies to define the unit of interaction testing and turns the homoeolog groups of an allopolyploid genome into one testable family with experiment-wide error calibration. Each homoeolog group of two, three or four copies is tested through its pairwise edges with three complementary genotype encodings combined into one omnibus P value, and error is controlled across the whole family by a parametric bootstrap minimum-P procedure with a structure-aware variance model. A companion workflow partitions additive variance by subgenome. We show that the interaction test is calibrated under the tested null models on the real genomes of four crops at family sizes from 80 to 17,404 groups, that it is powerful across interaction architectures and complements SNP-by-SNP scans, and that it detects homoeolog-group interactions for heading date in wheat, flowering time in rapeseed and fibre length in cotton. The same per-copy features show that homoeolog coupling across *B. napus* accessions has a weak but reproducible association with population-level homoeologous-exchange frequency. Complete interaction analyses of these panels take minutes, and the software is available with an interface for AI assistants.

## Results

### HomoeoGWAS turns homoeolog structure into a testable interaction family

Allopolyploid genomes carry their own map of candidate interactions: the homoeologous copies of each gene. HomoeoGWAS uses this map to organize interaction testing (Fig. 1). We applied it to hexaploid bread wheat (827 Watkins accessions; 2,143 A–B–D homoeolog triads passing a genotype-only design gate), tetraploid upland cotton (1,245 accessions; 3,757 A–D pairs), *Brassica napus* rapeseed (926 accessions; 17,404 A–C pairs) and tetraploid peanut (230 accessions; 379 A–B pairs) (Fig. 1a). The pairings were locally collinear for 95.0–99.9% of edges in wheat, cotton and rapeseed, against at most 17% under partners permuted within the same chromosome pair, and for all nine units detected below; the small peanut family was too sparse for this check to discriminate (Supplementary Table 1). The interaction workflow tests predefined homoeolog groups (Fig. 1c). A group of two, three or four copies contributes one, three or six pairwise edges; each edge is tested with three encodings of the two gene blocks (minor-allele burden product, first-principal-component product and a kernel-Hadamard term), and the encoding and edge P values are combined by the Cauchy combination into one group P value (group-omniB). All groups of a declared experiment form one family, whose family-wise error rate (FWER) is controlled by a parametric-bootstrap minimum-P procedure (bootstrap-minP) drawn from the fitted null model; in the primary null (smooth_pc4), sample-specific variance weights, estimated from squared whitened residuals as a smooth function of the top four genomic principal components, are applied to the fitted null covariance. Encoding and edge P values localize the evidence within a group. A companion additive workflow builds one genomic relationship matrix per subgenome, estimates their variance components jointly by multi-kernel restricted maximum likelihood (REML) and reports the proportion of phenotypic variance explained (PVE) by each subgenome with a leave-one-chromosome-out (LOCO) marker scan (Fig. 1b). Every run binds its inputs by SHA-256 and stops rather than returning partial results when a preflight check fails; the software runs from the command line or Python, with an optional interface for AI-assisted analysis (Fig. 1d).

**Fig. 1.**
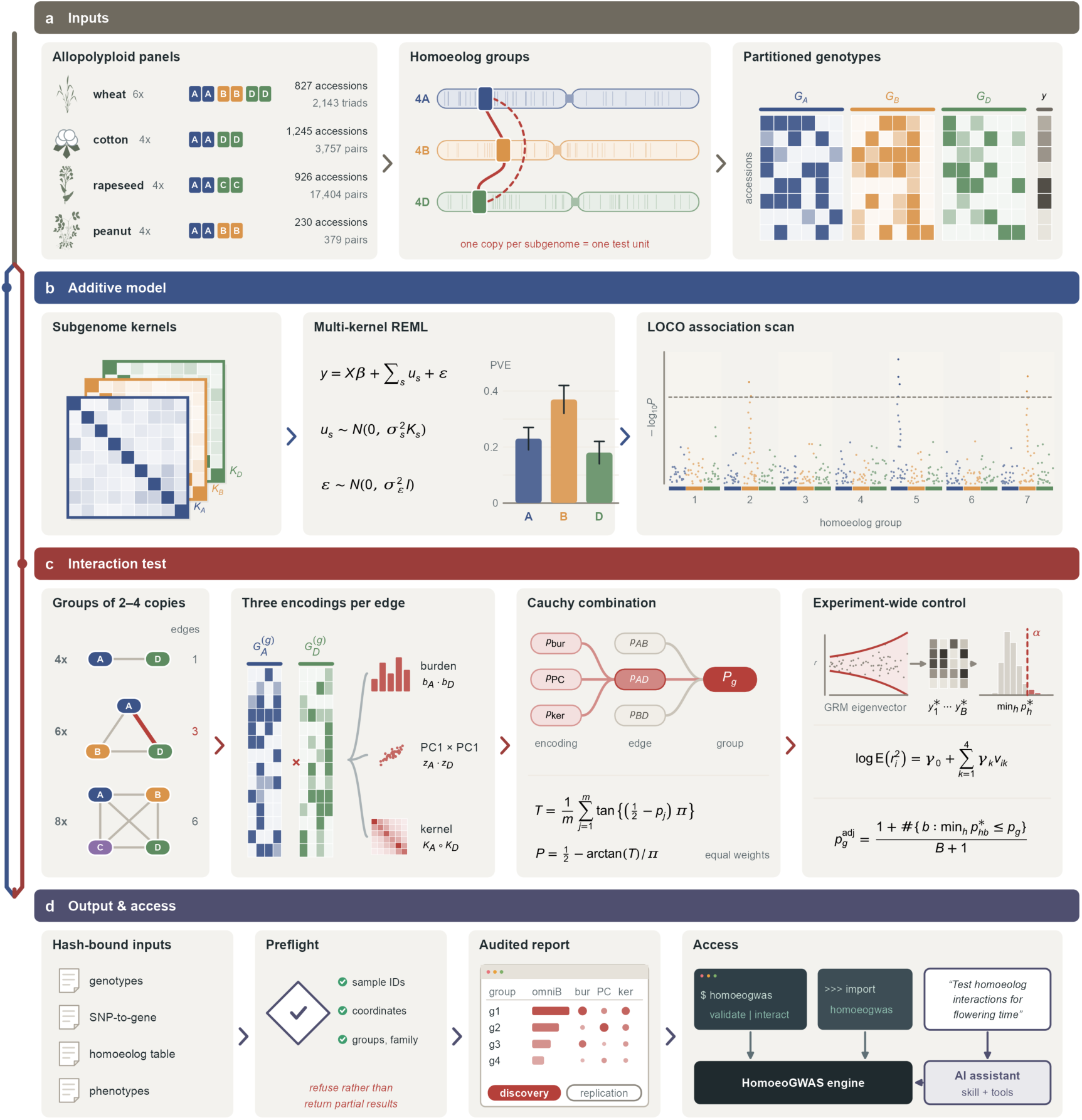
HomoeoGWAS workflow. Additive (blue track) and interaction (red track) analyses share the inputs and one output. **a**, Panels (subgenome composition; accessions; tested homoeolog groups) and a homoeolog group, one copy per subgenome. *G_s_*, genotypes of subgenome *s*; *y*, phenotype. **b**, Additive model: subgenome relationship matrices *K_s_*, multi-kernel REML, subgenome proportions of phenotypic variance explained (PVE) and a leave-one-chromosome-out (LOCO) scan (illustrative values). **c**, Interaction test. Groups of two, three or four copies (4x, 6x, 8x) give one, three or six edges; each edge is tested with the burden product (*b_A_* ⋅ *b_D_*), the PC1 product (*z_A_* ⋅ *z_D_*) and a low-rank kernel-Hadamard term (*K_A_* ∘ *K_D_*). Encoding P values are combined by the Cauchy statistic *T* into edge P values (*p_AB_*, *p_AD_*, *p_BD_*) and the group P value *P*_g_. Experiment-wide control: variance function (smooth_pc4) of the whitened residuals *r_i_* on genomic eigenvectors *v_ik_*, null phenotypes *y*_1_^∗^ … *y_B_*^∗^ and the distribution of family minima min_ℎ_*p*^∗^ with the family-wise level α. **d**, Hash-bound inputs, preflight checks, report and access routes.

### Experiment-wide error control is calibrated on four crop genomes

We first simulated null phenotypes on the real genotypes of the wheat, rapeseed and cotton panels, each with a family of 80 homoeolog groups, and analysed them with the complete pipeline, including per-phenotype REML refitting and bootstrap-minP (B = 2,000, α = 0.05; Fig. 2a; Supplementary Fig. 1; Supplementary Data 1, sheets S2-3 to S2-7). Under a kinship-preserving Gaussian null the empirical FWER was 3.9% in wheat (49 of 1,250; Wilson 95% confidence interval 3.0–5.1%) and 4.6% in cotton (57 of 1,250; 3.5–5.9%) with the earlier homoscedastic null, and 4.5% in rapeseed (56 of 1,250; 3.5–5.8%) with smooth_pc4. Nulls carrying an additive or a PC1 main effect gave 1.4–4.4% (500 phenotypes each).

**Fig. 2.**
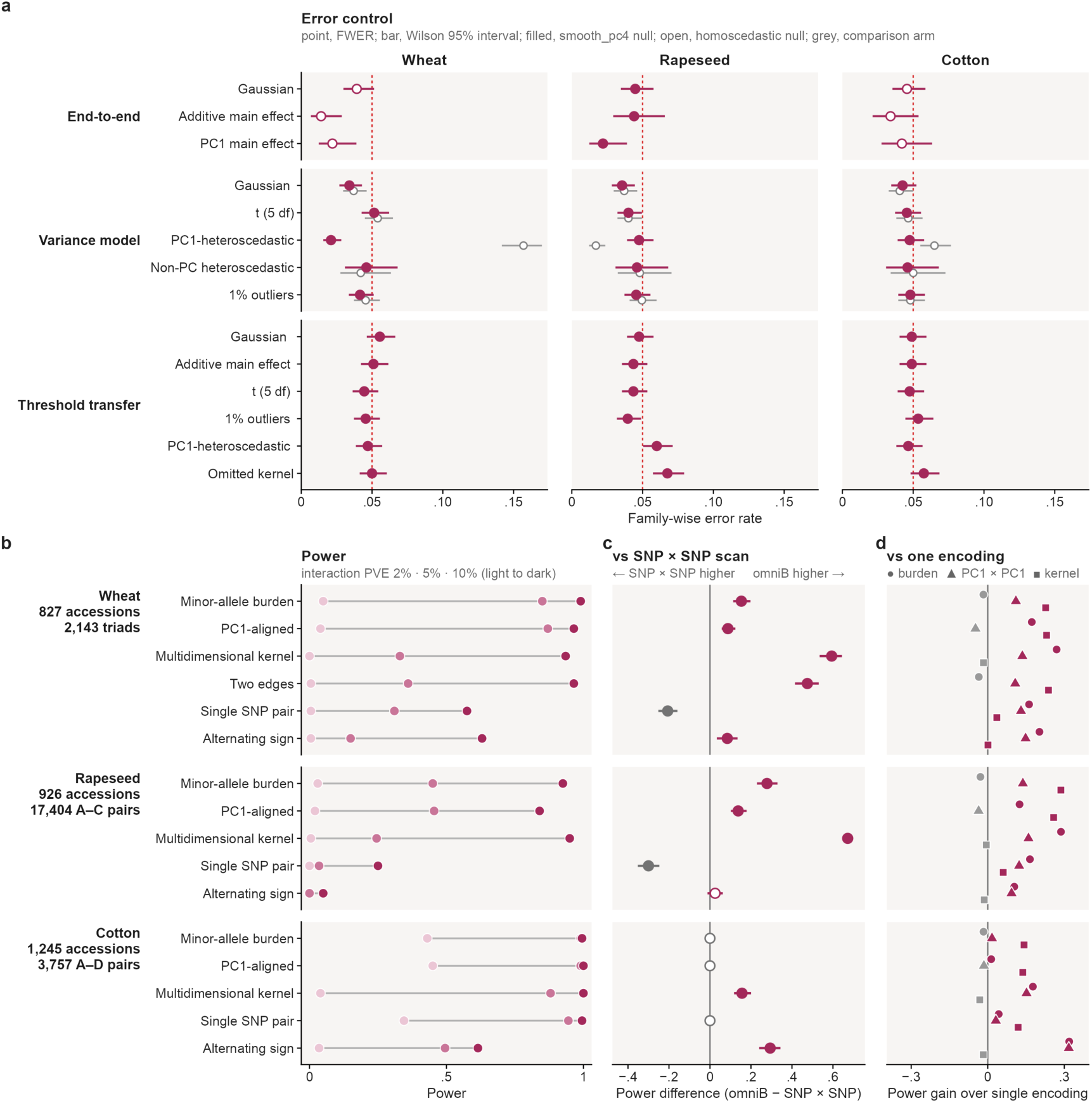
HomoeoGWAS controls family-wise error and gains power across three crop genomes. **a**, Family-wise error rate (FWER; α = 0.05) in 80-group families. End-to-end: complete pipeline under Gaussian (1,250 phenotypes), additive and PC1 main-effect nulls (500 each); open, homoscedastic null; filled, smooth_pc4. Variance model: five noise models; magenta, smooth_pc4; grey, homoscedastic null. Threshold transfer: fixed whitening, no bootstrap, 2,000 held-out phenotypes per layer. Bars, Wilson 95% intervals; dashed line, 0.05. **b**, Power at interaction PVE 2%, 5% and 10% (light to dark), one planted interaction per phenotype in the complete family; 200 phenotypes per point. **c**, Power difference between group-omniB and a regenie SNP × SNP scan (320 phenotypes per cell; PVE 5%; matched empirical error); bars, Newcombe 95% intervals; filled, interval excludes 0. **d**, Power gain of group-omniB over each single encoding, mean of six conditional cells (PVE 2%, 5% and 10% × one or four causal groups); points beyond ±0.35 drawn at the axis limit; magenta, omniB higher.

When residual variance followed population structure, the homoscedastic bootstrap null over-rejected in wheat (FWER 0.157) and cotton (0.065), although not in rapeseed (0.017; 2,000 phenotypes per cell). We therefore adopted smooth_pc4, which applies structure-aware sample weights to the fitted null covariance, as the primary null and validated it on fresh phenotypes (Methods). Across five noise models (Gaussian, Student-t, 1% contamination at 6 s.d., and PC1-aligned and non-PC heteroscedasticity) and three panels, smooth_pc4 held the FWER between 0.021 and 0.052, with every Wilson upper bound at or below 0.068 (Fig. 2a; Supplementary Fig. 1b). In a separate threshold-transfer diagnostic with fixed whitening and no bootstrap, learned thresholds also transferred to independent held-out phenotypes (3.8–6.2% across five null layers and all statistics, with an omitted-kernel diagnostic reported separately; Supplementary Fig. 1a).

Finally, we tested calibration at the scale of the real families. In prespecified experiments, null phenotypes combining a polygenic component (generator weight 0.30) with residual variance aligned with the first two genotype principal components (weight 0.70) were reanalysed over each complete family with the delivered pipeline (smooth_pc4, B = 2,000; 640 phenotypes per panel; Supplementary Table 2). Any-group rejection was 1.4% for the 2,143 wheat triads (one-sided 95% upper bound 2.4%), 3.1% for the 17,404 rapeseed pairs (4.5%), 1.1% for the 3,757 cotton pairs (2.0%) and 3.1% for the 379 peanut pairs (4.5%), with no failed analysis. Families of 500 and 2,000 groups gave 1.2–5.8% (Supplementary Fig. 2). Error control was therefore maintained from 80 to 17,404 groups under the tested global-null models.

### The omnibus improves detection of distributed interactions and complements SNP-by-SNP scans

To estimate the power a user would obtain, we planted one pairwise interaction per phenotype in a unit drawn uniformly from the complete family, on the same background, and reanalysed each phenotype with the delivered pipeline (Fig. 2b; Supplementary Data 1, sheet S2-2). At a nominal PVE of 10%, defined on the untransformed simulated phenotype, power was at least 0.84 in 11 of the 16 panel-by-architecture combinations (six architectures in wheat, five in each two-copy panel; 200 phenotypes each). The hardest architectures were an alternating-sign signal that cancels an unsigned burden (wheat 0.63, cotton 0.62, rapeseed 0.05) and a single causal SNP pair in wheat and rapeseed (0.58 and 0.25), and power was lowest in rapeseed, whose family is by far the largest.

The three encodings were complementary. Each single encoding failed badly on some architecture, such as the burden under alternating signs, whereas group-omniB fell more than 0.02 below a single encoding in only 32 of 288 cell-by-encoding comparisons in an 80-group conditional grid (largest deficit 0.17; Fig. 2d; Supplementary Fig. 3; Supplementary Data 1, sheet S2-9). The omnibus thus reduces the need to know the interaction architecture in advance.

Group-omniB was also more powerful than a SNP-by-SNP scan when interaction was distributed over a group. We compared it with a regenie SNP × SNP product-term scan aggregated to the group by minimum P, on the same 320 phenotypes per cell in 80-group families (5% PVE, one causal group), with both methods calibrated to the same empirical error under Gaussian nulls (held-out FWER 4.8–5.7%; Fig. 2c; Supplementary Fig. 4b; Supplementary Data 1, sheet S2-10). Group-omniB detected significantly more in 10 cells, mostly where the interaction spanned several genotype dimensions or edges (wheat multidimensional kernel, 0.82 versus 0.22); four cells were ties, and the SNP × SNP scan was more powerful in the two cells where one SNP pair carried the interaction (rapeseed, 0.59 versus 0.89). The two approaches are therefore complementary, with the group test favoured when interaction is distributed over a homoeolog group.

### Detections localize a homoeolog group and its region

We next asked what a significant group identifies (Supplementary Figs. 3 and 4a; Supplementary Data 1, sheets S2-8 and S2-11). In the whole-family power simulations, rejections of non-causal groups fell mostly on the chromosomes carrying the causal unit’s copies, consistent with linkage, and occurred on other chromosomes in 2.1% of replicates; in cells with power of at least 0.80 the causal unit ranked first in 66–98% of replicates. Two controls examined specificity in the 80-group conditional grid at 10% PVE. Strong additive effects concentrated in one group, without interaction, raised the family-wise false-rejection rate to 8.0%, 9.0% and 12.8% in wheat, cotton and rapeseed when phenotypes were scored with a fixed whitening matrix on the untransformed scale and without the bootstrap. These rejections fell almost entirely on groups other than the one carrying the additive effects (0 of 42 rejected groups in wheat, 0 of 46 in cotton and 2 of 73 in rapeseed). With per-phenotype REML on the transformed phenotype, as in the delivered pipeline, a rescoring of 200 stored phenotypes per panel gave 6.0% in wheat (95% interval 3.5–10.2%) and 8.5% in cotton (5.4–13.2%), against 9.5% and 10.5% with fixed whitening (Supplementary Fig. 4c). A strong additive effect concentrated in one homoeolog group can therefore inflate the family-wise error above the nominal level, whereas genome-wide additive or PC1 main effects (1.4–4.4%, end-to-end calibration) and planted additive variants of up to 20% PVE outside the tested gene blocks (1.1–5.2%, whole families; leakage control below) did not; the conditions are compared in Methods. An interaction between one copy of a group and a gene outside it led to any-group rejection in 12.2–23.2% of replicates. Under the mispaired control, 22% and 20% of rejected groups in wheat and cotton were the group containing the reference copy and 16% and 18% the group of the external partner gene. A significant group therefore reports interaction evidence for the group and its genomic neighbourhood, and identifying the interacting partner requires follow-up of the group.

### Subgenome-stratified scans recover known loci and partition phenotypic variance

The additive workflow recovered known loci in each panel (Fig. 3a; Supplementary Table 3). For cotton fibre length (n = 1,245; genomic inflation factor 1.11), the scan recapitulated all four loci previously reported for the same accessions at genome-wide (Bonferroni) significance. For wheat heading date (days to ear emergence; n = 827; environment-adjusted line means with records beyond 6 s.d. of the trial mean set to missing), 3 of the 29 loci reported for the Watkins panel had a genome-wide significant SNP within ±5 Mb, against 0.18 expected (P = 0.002; genomic inflation factor 1.39), and Vrn-B1 and Vrn-D1 reached genome-wide significance in the 2006 trial. For rapeseed flowering time (n = 926; genomic inflation factor 1.12), 3 of 7 known flowering loci were recovered at P < 10⁻⁵, against 0.60 expected (P = 0.016); BnaFLC.A10 and the BnaFLC.A03b/FRI region approached but did not reach Bonferroni significance (P = 9.7 × 10⁻⁸ and 6.1 × 10⁻⁸; threshold 4.95 × 10⁻⁸). In peanut (n = 230), hundred-seed weight had two and seed length one SNP at Bonferroni significance, and AhARF2-2 was recovered at P = 1.8 × 10⁻⁶; GEMMA detected at least as many significant SNPs on the same data.

**Fig. 3.**
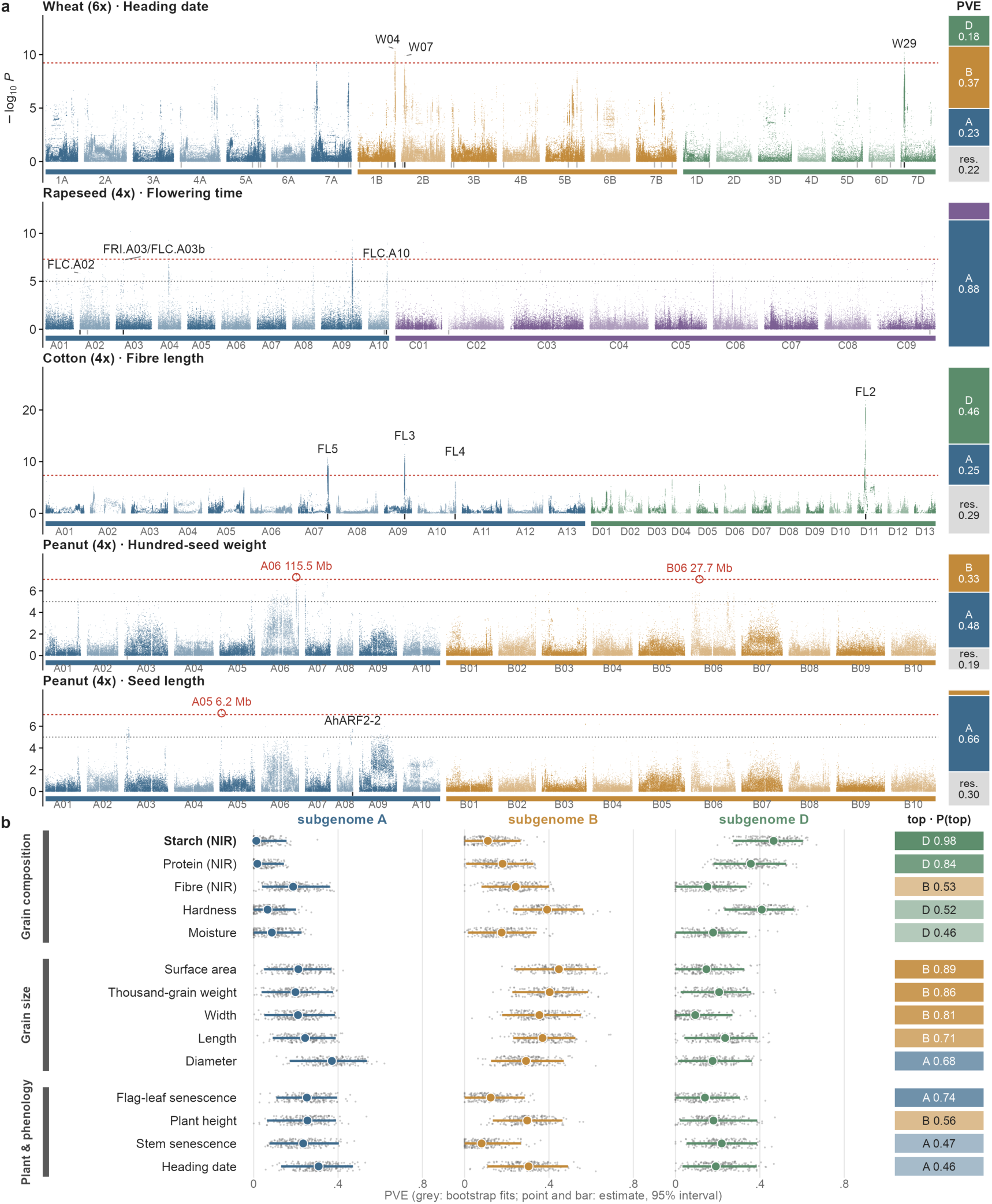
HomoeoGWAS subgenome-stratified scans and subgenome partitioning of phenotypic variance. **a**, Report cards for wheat heading date (n = 827), rapeseed flowering time (n = 926), cotton fibre length (n = 1,245) and peanut seed traits (n = 230). Manhattan plots, LOCO scans coloured by subgenome; red dashed line, Bonferroni threshold; grey dotted line, P = 10⁻⁵; red circles, Bonferroni-significant SNPs (peanut); ticks below the axis, known loci (black, recovered; grey, not recovered). Bars at right, PVE of each subgenome and the residual. **b**, Subgenome PVE of 14 wheat traits: grey points, 200 bootstrap fits; coloured point and bar, estimate and 95% interval; right, subgenome with the largest PVE and its bootstrap probability; bold, trait passing the multistart stability gate.

**Fig. 4.**
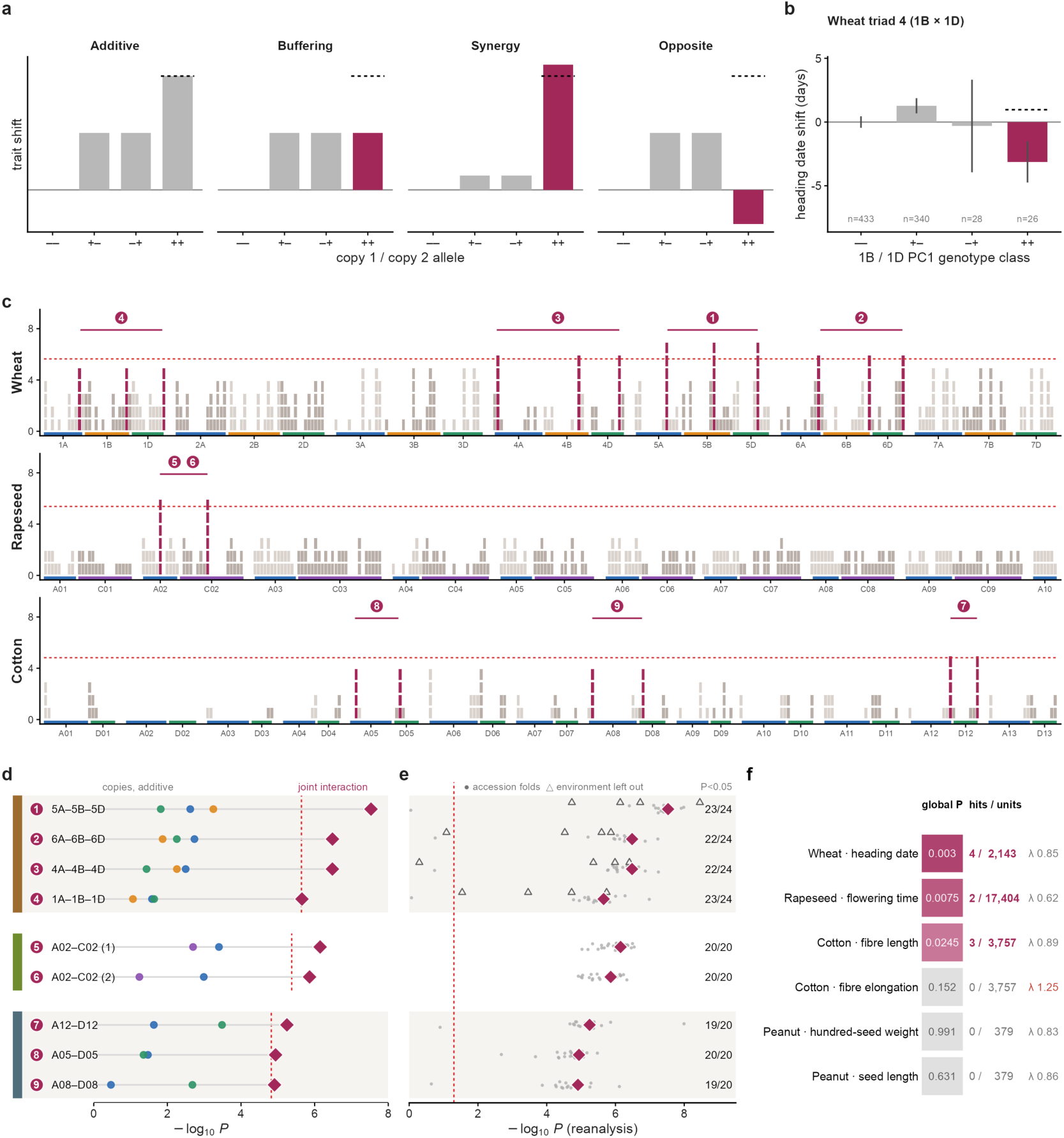
Homoeolog-interaction scans in four allopolyploid crops. **a**, Illustrative non-additive patterns of two copies; dashed line, additive expectation; magenta, double combination. **b**, Wheat triad 4 (1B × 1D): raw (unadjusted) mean heading date of PC1-defined genotype classes relative to –/– (two-means classes of the gene PC1 score, sign fixed so that the largest-magnitude score is positive; –, lower; +, higher); error bars, 95% t interval of the class mean; n, accessions per class; dashed line, additive expectation. **c**, Interaction scans of wheat (2,143 triads), rapeseed (17,404 pairs) and cotton (3,757 pairs). Each column is a genome window whose stacked squares give the −log10 P of its strongest group (one square per unit; windows with −log10 P ≥ 1.5 or a significant group); magenta, groups significant at 5% FWER, copies joined by a bar; red dashed line, experiment-wide threshold; 1–9, significant units. **d**, Strongest additive −log10 P of each copy (coloured by subgenome) and interaction −log10 P of the group (diamond); dashed line, interaction threshold. **e**, Nominal P of each unit with the formal features fixed in 20 accession-deletion folds (grey) and leave-one-environment reanalyses (triangles, wheat); diamond, full data; dashed line, P = 0.05; k/N, reanalyses with P < 0.05 (N = 20; 24 in wheat). **f**, Global bootstrap-minP P (tile; grey, not significant), significant over tested units and interaction-scan inflation factor λ (red, λ > 1.15) for each experiment. Units 1–9: wheat chromosome 5, 6, 4 and 1 triads; rapeseed A02–C02 pairs; cotton A12–D12, A05–D05 and A08–D08 pairs.

Subgenome partitioning ranked the subgenomes by the variance they explain. Across 14 wheat traits with 200 bootstrap fits each (multistart-audited for four traits), NIR grain starch was the only trait that passed the prespecified stability gate (Fig. 3b; Supplementary Table 4). The D subgenome explained 0.465 of starch variance (95% bootstrap interval 0.274–0.604) and ranked first in 98.0% of bootstrap fits; its share of the genetic variance (0.79) lay at the 96.8th percentile of random chromosome groupings. Point estimates for the application traits are given in Supplementary Fig. 5.

### Homoeolog-interaction scans in four allopolyploids

We applied the interaction workflow to each application trait with one frozen family and one bootstrap-minP correction per experiment (smooth_pc4, B = 2,000; Fig. 4f; Supplementary Table 5). For wheat the phenotype was the environment-adjusted heading date, inverse-normal transformed within the pipeline. Four of 2,143 wheat triads (global P = 0.003, the bootstrap-minP-adjusted P value of the most significant group), two of 17,404 rapeseed pairs forming one locus (0.0075) and three of 3,757 cotton pairs for fibre length (0.0245) were significant at 5% FWER (units 1–9 in Fig. 4c–e). Cotton fibre elongation (global P = 0.152) and, in the small peanut panel (230 accessions), hundred-seed weight (0.99) and seed length (0.63) yielded no significant group. The fibre-elongation scan had a genomic inflation factor of 1.25 and is flagged for calibration review.

The detected groups showed no strong nearby additive associations and were not unusually aligned with the leading axes of population structure. The copy-resolved additive signal of each unit is shown in Fig. 4d; no unit had an additive SNP with P < 10⁻⁵ within ±1 Mb, and their alignment with population structure fell within the matched background (10th–66th percentiles; Supplementary Table 6). In a prespecified control, additive variants planted outside every tested gene block and correlated with the detected interaction features left the experiment-wide rejection rate within the 7.5% tolerance in all nine panel-by-strength cells (1.1–5.2%), and no detected unit was withdrawn (Supplementary Table 7). With the formal features held fixed, each unit kept nominal support (P < 0.05) in at least 19 of 20 accession-deletion folds, and in wheat two triads kept support in all four leave-one-environment reanalyses and two in three of four (Fig. 4e; Supplementary Table 8).

Conditioning on regional genotype principal components, a post-selection robustness analysis, separated two tiers of evidence. Five groups at four loci retained their evidence: the wheat chromosome 5 and 6 triads (units 1 and 2), the two rapeseed A02 × C02 pairs at one locus (units 5 and 6) and the cotton A12 × D12 pair (unit 7). The wheat chromosome 4 triad (unit 3) was uncertain, and the wheat chromosome 1 triad (unit 4) and the cotton A05 × D05 and A08 × D08 pairs (units 8 and 9) were not robust; we treat these as candidate regional signals. With the homoscedastic null, six further wheat triads were rejected; they are listed as a sensitivity tier and not claimed, because that null over-rejects under structure-aligned heteroscedasticity in this panel (Supplementary Tables 9 and 10).

Among the robust signals, unit 1 is the strongest wheat detection (adjusted P = 0.003) and carries orthology and domain support for a ZnF-UBP ubiquitin-specific protease^28^ (rice OsUBP2), making it the lead for follow-up. Fig. 4b illustrates the allelic pattern the test targets (Fig. 4a) with unit 4, a regional candidate and the only detected triad with at least 26 accessions in every genotype class: in raw phenotype means not adjusted for relatedness or population structure, accessions in the higher PC1-defined genotype class at both the 1B and 1D copies (n = 26) headed about 4 days earlier than expected from the two single-copy classes (difference of differences −4.1 days; bootstrap 95% interval −8.6 to −0.7). Prespecified three-copy conditional models of the same trait found no experiment-wide A × B × D effect (Supplementary Table 11).

### Homoeolog coupling is associated with homoeologous exchange in rapeseed

The per-copy features used by the interaction test also reveal how homoeologs co-vary across accessions, independently of any trait. For each edge we defined homoeolog coupling *r_P_* as the absolute correlation across accessions between the two copies’ gene PC1 features after adjustment for ten genotype principal components; because each accession contributes one product term, coupling decomposes by accession, and its null distribution re-pairs each copy with a non-homoeologous copy on another chromosome (Fig. 5a). Rapeseed coupling was highly reproducible: its full-panel reliability lower bound of 0.704 exceeded the reliability threshold of 0.138 (median half-panel correlation 0.567; Fig. 5f).

**Fig. 5.**
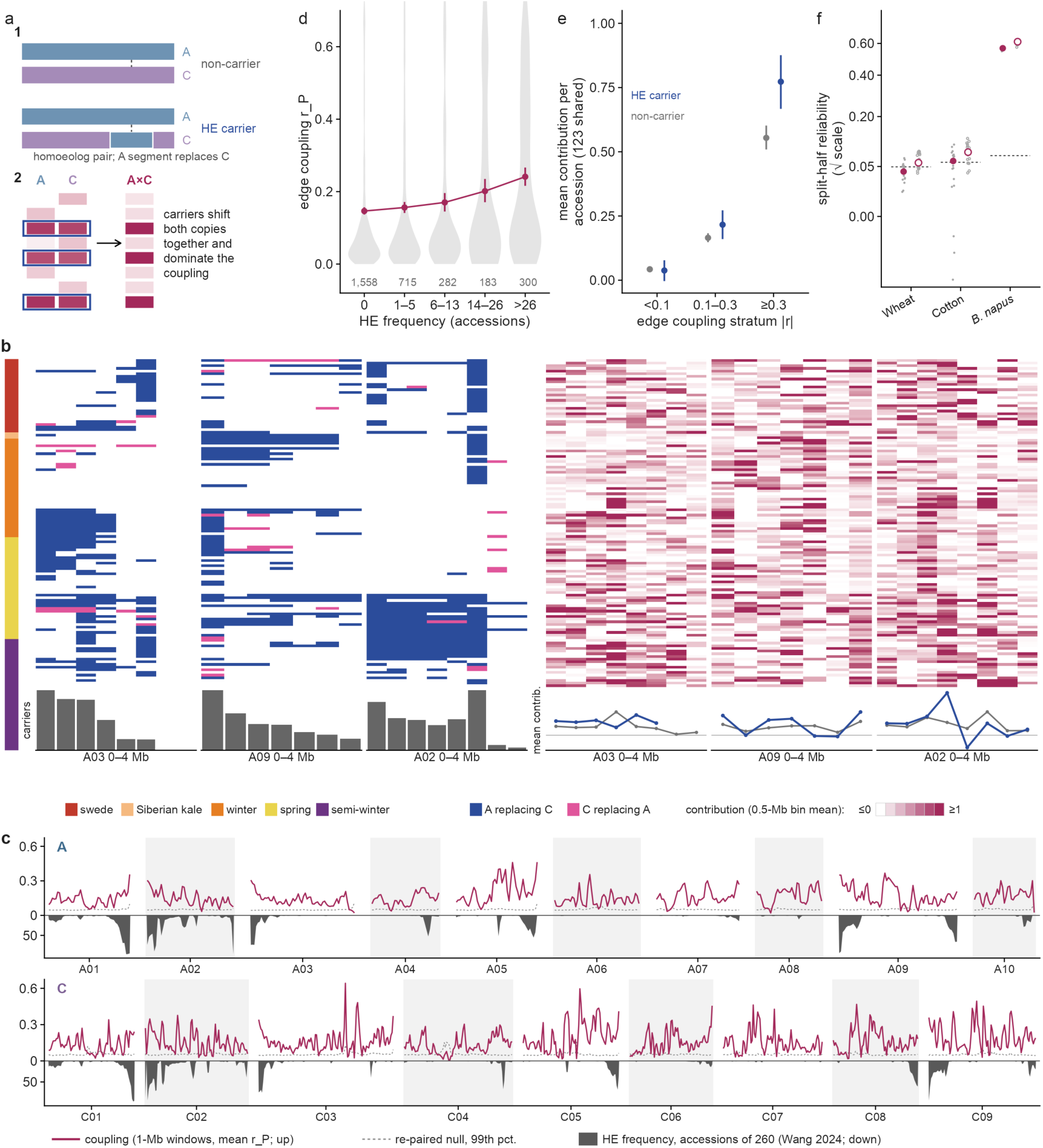
Homoeolog coupling is associated with homoeologous-exchange frequency in B. napus. **a**, Schematic of a homoeologous exchange (HE) and of coupling *r_P_*, the absolute correlation across accessions of the structure-adjusted gene-PC1 features of the A and C copies, which decomposes into per-accession contributions. **b**, Three HE hotspots: on A03, A09 and A02, the 2-Mb A-copy window with the highest published HE frequency^29^ among windows with ≥ 8 mapped edges, shown within 0–4 Mb. Rows, the 123 accessions shared with Wang et al., ordered by ecotype and number of exchanges. Left, HE calls (blue, A replacing C; pink, C replacing A); right, mean per-accession contribution in 0.5-Mb bins (white ≤ 0 to magenta ≥ 1); bottom, carriers per edge and mean contribution of carriers (blue) and non-carriers (grey). **c**, Coupling in 1-Mb windows (magenta, upwards) with the 99th percentile of the re-paired null (dashed) and HE frequency (grey, downwards; accessions of 260), A and C subgenomes. **d**, Edge coupling by HE frequency (0, 1–5, 6–13, 14–26 and > 26 carriers; 3,038 edges): grey, distribution; magenta, mean and 95% edge-bootstrap interval. **e**, Mean contribution of carriers (blue) and non-carriers (grey) by coupling stratum (post hoc strata; 95% accession-bootstrap intervals). **f**, Split-half reliability of edge coupling in 20 splits (grey; magenta, median; filled, *r_P_*; open, kinship-whitened); dashed line, half-panel reliability needed for 80% power at ρ = 0.13.

Rapeseed coupling rose with the frequency of homoeologous exchange (HE) reported for 260 *B. napus* accessions^29^. Across the 3,038 of 17,404 edges that map one-to-one to that study’s anchors, the association was weak but reproducible (Spearman ρ = 0.136; within-chromosome circular-shift permutation P ≤ 5 × 10⁻⁴), with mean *r_P_* rising from 0.146 for edges without carriers to 0.241 for edges with more than 26 (Fig. 5c,d; Supplementary Fig. 6). It persisted after post hoc adjustment for SNP count, minor-allele frequency and heterozygosity (partial ρ = 0.133) and was positive in 9 of 10 A chromosomes, consistent with a contribution of recurrent homoeologous exchange to homoeolog coupling across accessions. Filtering heterozygous calls removed most of the genome-wide coupling excess over re-pairing, whereas the association with exchange frequency was unchanged after adjustment for per-edge heterozygosity (ρ = 0.137).

Accessions carrying reported exchanges contributed disproportionately to the aggregate coupling statistic at strongly coupled edges. In the 123 accessions shared with that study (identical names and sequencing runs), the mean per-accession contribution across the mapped accession–edge pairs was higher for carriers than for non-carriers (0.307 versus 0.162), exceeding a null that reassigns whole HE profiles within ecotypes (P ≤ 0.001; Fig. 5b,e). In post hoc strata the carrier excess was concentrated at strongly coupled edges (0.176 at *r_P_* ≥ 0.3, P = 0.0033, the smallest value attainable with 300 permutations; 0.036 at 0.1–0.3, P = 0.07; −0.011 below 0.1, P = 0.77). At strongly coupled edges (*r_P_* ≥ 0.3), therefore, the accessions that carry an exchange contribute disproportionately to the coupling; this is a population-level pattern rather than a classifier of individual carriers (area under the curve 0.519 against 0.518 for permuted profiles; Supplementary Fig. 7), and HE calls and genotypes share sequencing data.

Wheat and cotton showed weaker coupling with lower edge-level reliability (full-panel lower bounds 0.052 and 0.050 against 0.094 and 0.111; a kinship-whitened cotton estimator passed narrowly; Fig. 5f; Supplementary Table 12). In cotton, coupling exceeded the re-pairing null (mean 0.039 versus 0.034; P = 0.001, also after kinship whitening) and passed the prespecified within-cultivar-group test marginally (one-sided P = 0.042); in a post hoc diagnostic it disappeared when principal components were re-estimated within groups, so it varies with finer population substructure (Supplementary Fig. 8). In wheat, the genome-wide excess was small (0.00105; one-sided 95% upper bound 0.0021), and associations with introgression, copy number and expression co-variation were not separable from variant density or absent (Supplementary Fig. 6). These panels thus bound the aggregate coupling signal at the precision their edge-level measurements allow.

### Subgenome-aware and experiment-wide analyses run in minutes

We measured wall time and peak memory on two nodes, each with two 64-core AMD EPYC 7742 processors, as medians of three repeats (four for mvMAPIT), with declared thread budgets for every tool (Fig. 6; Supplementary Table 13). On wheat with 83.3 million markers and 827 accessions at 64 threads, the HomoeoGWAS scan with A, B and D kernels and LOCO took 17 min, REGENIE 20 min and GEMMA, with one pooled kinship, 10.5 h (Fig. 6a); wall time grew with the number of accessions up to 5,000 simulated genotypes at 10 million markers (Fig. 6b). On the 10-million-marker wheat scan, three subgenome kernels with LOCO cost about 18% more than one kernel with LOCO (Fig. 6d), and wall time fell 5.5-fold from 1 to 256 threads with little gain beyond 32 (Fig. 6e). At 5,000 simulated genotypes HomoeoGWAS used more memory than GEMMA (52 versus 6.5 GB; Fig. 6f).

**Fig. 6.**
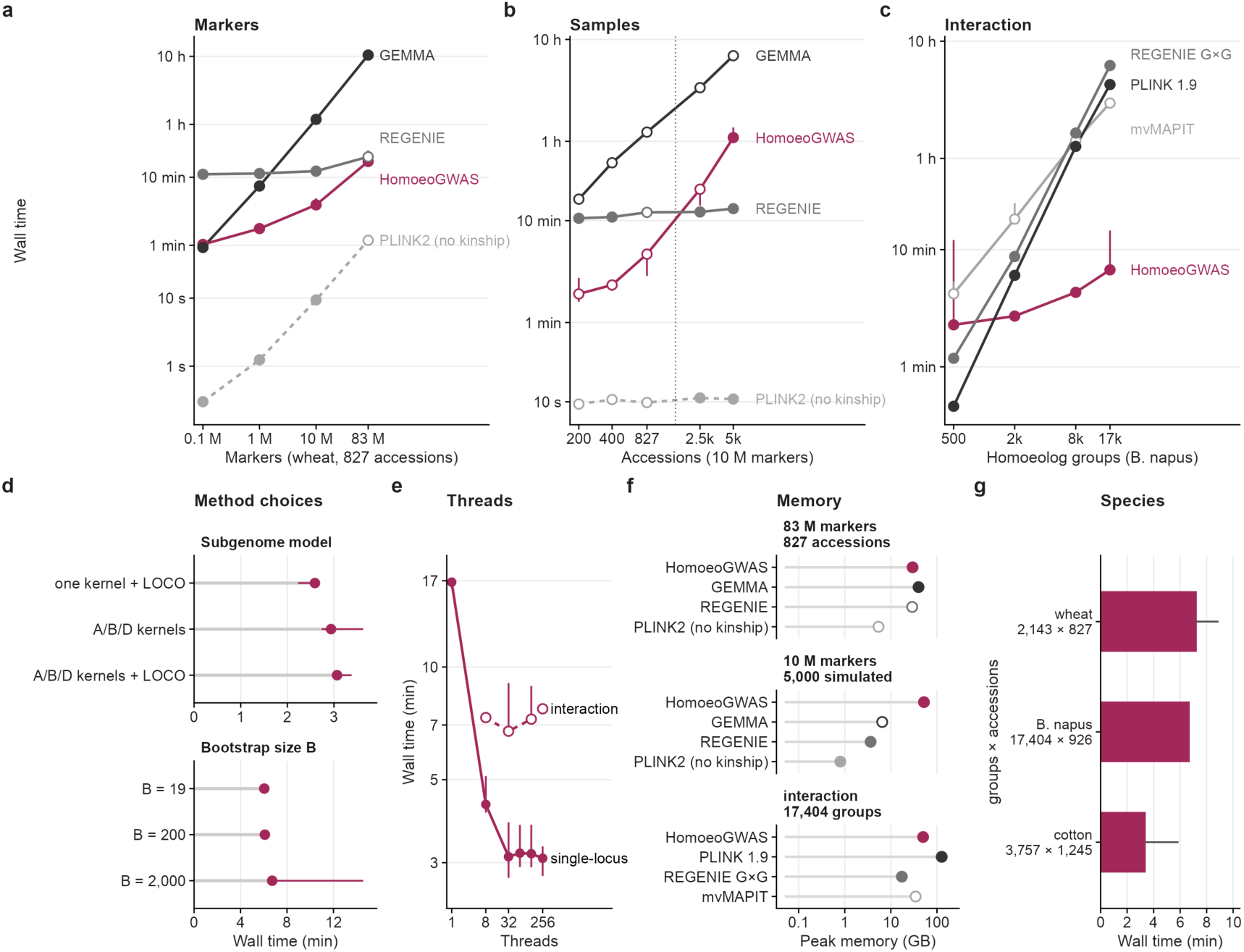
Computational cost of subgenome-aware scans and experiment-wide homoeolog-interaction analysis. Points, median of three repeats (four for mvMAPIT); bars, range; open symbols, repeats overlapping another heavy job. **a**, Single-locus scan of wheat (827 accessions; 64 threads) against marker number: HomoeoGWAS (A, B and D kernels with LOCO), GEMMA (one kinship), REGENIE and PLINK2 (no kinship; computational lower bound). **b**, The same against the number of accessions at 10 M markers (2,500 and 5,000 simulated, right of the dotted line). **c**, Complete interaction analysis (B = 2,000; 128 workers) against the number of B. napus groups, with tools that test other hypotheses: PLINK 1.9 (all SNP pairs), REGENIE G×G (within-group SNP pairs) and mvMAPIT (marginal epistasis). **d**, Cost of subgenome kernels and LOCO (top) and of the number of bootstrap replicates B (bottom). **e**, Thread scaling of the single-locus scan and the interaction analysis. **f**, Peak memory in three workloads. **g**, Complete interaction analysis per species (groups × accessions). Tool settings, Supplementary Table 13.

With bootstrap-minP (B = 2,000) and 128 workers, the complete interaction analysis took 7.3 min for wheat (2,143 triads), 6.7 min for rapeseed (17,404 pairs) and 3.4 min for cotton (3,757 pairs) (Fig. 6g). Experiment-wide calibration added about 11% over B = 19 (Fig. 6d), and the wheat analysis took 6.7–7.7 min across 8–256 workers (Fig. 6e). At 17,404 rapeseed groups and 64 declared threads, tools that test other hypotheses took 3.0 h (mvMAPIT, marginal epistasis, on the node used for single-locus scans), 4.3 h (PLINK 1.9, all SNP pairs) and 6.2 h (REGENIE, within-group SNP pairs) (Fig. 6c); these figures compare cost under each tool’s configuration, and the highlighted REGENIE single-locus and mvMAPIT measurements include repeats flagged for interference from other jobs.

The software is available as a command-line tool, a Python library and an interface for AI assistants, whose evaluation on 30 frozen analysis tasks is reported in Supplementary Fig. 9 and Supplementary Table 14.

## Discussion

HomoeoGWAS defines the unit of genetic-interaction testing by evolutionary homology: the group of homoeologous gene copies that allopolyploidy created. Treating all such groups in a genome as one family turns homoeolog interaction into a genome-wide association analysis with experiment-wide error calibration. HomoeoGWAS adapts established P-value combination and resampling procedures^30,31^ to this biologically defined unit, which makes genome-wide homoeolog interaction mapping feasible while accounting for multiplicity across the complete family of eligible groups; each group is evaluated with encodings that capture dosage-like, haplotype-like and multidimensional interaction. This reorganisation has three consequences shown here. First, the family becomes small enough to calibrate by resampling from the fitted null model: calibration held under Gaussian, heavy-tailed, contaminated and structure-aligned heteroscedastic nulls in wheat, cotton and rapeseed, and under a structure-aligned global null over the complete families of all four crops, up to the 17,404 A–C pairs of rapeseed. Second, the omnibus is powerful without prior knowledge of the interaction architecture, because its encodings are complementary: each single encoding failed on some architecture that the combination detected. Third, the analysis is fast: complete interaction analyses of the benchmark panels took 3.4–7.3 min with 2,000 bootstrap replicates and 128 workers, whereas tools testing all SNP pairs or marginal epistasis took hours for the same groups.

The comparison with a SNP-by-SNP scan shows when a group test is the right tool. In 80-group families with thresholds calibrated empirically under Gaussian nulls, group-omniB detected substantially more when simulated interaction was spread over several genotype dimensions or edges of a homoeolog group, and the pairwise scan was more powerful when a single SNP pair carried the signal. The two approaches are complementary: a group test aggregates evidence that a pairwise scan dilutes across many tests, and a pairwise scan resolves a concentrated signal that a group test averages. A natural workflow is therefore a group scan across the genome, followed by SNP-level analysis inside significant groups.

The crop applications show what the test finds in practice. Nine units at eight loci were significant at 5% experiment-wide error for heading date in wheat, flowering time in rapeseed and fibre length in cotton. No additive SNP reached P < 10⁻⁵ within 1 Mb of the detected units, none of the nine units met the prespecified withdrawal criterion in the additive-leakage simulations, and each kept nominal support in at least 19 of 20 accession-deletion reanalyses. Conditioning on regional genotype structure separated five robust groups at four loci — the wheat chromosome 5 and 6 triads, the rapeseed A02 × C02 locus and the cotton A12 × D12 pair — from an uncertain wheat chromosome 4 triad and three regional candidates (the wheat chromosome 1 triad and the cotton A05 × D05 and A08 × D08 pairs). The wheat chromosome 5 triad, the strongest detection, carries a candidate ZnF-UBP ubiquitin-specific protease supported by orthology and protein-domain evidence, providing a target for future homoeolog-specific functional validation.

Homoeolog coupling adds a genome-level view that interaction scans alone do not give. In *B. napus*, coupling between A and C copies was highly reproducible, rose with the population frequency of homoeologous exchange and, at strongly coupled edges (*r_P_* ≥ 0.3), was carried disproportionately by the accessions that carry an exchange. This association supports a population-level relationship between homoeolog coupling and reported exchange frequency, rather than individual-level prediction of carriers, and the strongly coupled edges are candidates for studying how exchange shapes trait variation^29,32^. In wheat and cotton, coupling was weaker and its edge-level measurement less precise, so these panels bound the aggregate signal rather than resolving it locus by locus.

These results define how HomoeoGWAS detections are read. Calibration was demonstrated under the null generators tested, including heteroscedasticity in which sample-specific variance weights vary smoothly with the leading genomic principal components; it does not extend to a strong additive effect concentrated in one homoeolog group, and strong control of the family-wise error rate when some groups carry true interactions was not established. A significant group locates interaction evidence in a homoeolog group and its genomic neighbourhood, and identifying the interacting partner and the causal genes calls for follow-up such as allele-specific expression, near-isogenic lines or homoeolog-specific mutants and editing^33,34^. Independent-population replication will be important for assessing the transferability of these signals. The framework applies to allopolyploids with divergent subgenomes and established homoeolog maps; autopolyploids, mixed-ploidy panels and direct higher-order interaction coefficients are outside its scope.

Allopolyploid crops are being resequenced at scale^15,35–39^, and their homoeolog maps improve with each new assembly. HomoeoGWAS provides a calibrated, fast and reproducible way to ask of these resources a question that single-marker association cannot: which homoeolog groups act together to shape a trait.

## Methods

### Panels, genotypes and homoeolog groups

We analysed four allopolyploid diversity panels. Wheat (*Triticum aestivum*, AABBDD): 827 phenotyped accessions of the Watkins landrace collection (ENA PRJEB71453)^36^, with variants on IWGSC RefSeq v1.0 coordinates and v1.1 gene models^1^. Cotton (*Gossypium hirsutum*, AADD): 1,245 accessions of CottonGVD^37^ on the Gh_CRI v1 assembly. Rapeseed (*Brassica napus*, AACC): 926 accessions of BnaGVD^38^ with flowering-time records^40^ on the Darmor-bzh v4.1 assembly^3^. Peanut (*Arachis hypogaea*, AABB): 230 accessions of PeanutPan^39^ on the NDH108 assembly. Biallelic SNPs were assigned to subgenomes by chromosome of origin (wheat A/B/D chromosomes; cotton A01–A13/D01–D13; rapeseed A01–A10/C01–C09; peanut Chr01–10 = A, Chr11–20 = B) and stored as one PLINK file set per subgenome. Variant filtering was panel-specific: wheat single-locus scans used 83,296,047 SNPs with minor-allele frequency (MAF) ≥ 0.01 and call rate ≥ 0.90; cotton inputs were prefiltered at MAF ≥ 0.10 and scanned at MAF ≥ 0.05 (1,105,457 SNPs); rapeseed inputs were biallelic SNPs with MAF ≥ 0.05 and missingness ≤ 0.20 (1,009,546 SNPs); peanut inputs were 587,385 SNPs with MAF ≥ 0.05. The interaction features applied a further MAF gate of 0.01 within the analysed samples.

Homoeolog groups contain one gene per subgenome. Wheat triads were taken from the high-confidence 1:1:1 triads of Ramírez-González et al.^8^. Of the 17,036 high-confidence triads, 12,003 had at least three SNPs with minor allele frequency ≥ 0.01 in every copy in the Watkins panel; 11,920 of these had an estimable design, and a genotype-only design gate that used no phenotype retained the 2,143 triads with an effective sample size of at least 20 for the conditional target, a maximum single-accession information fraction of at most 0.20, a target residual ratio of at least 0.10 and at least five SNPs in every copy. For cotton (3,757 A–D pairs), rapeseed (17,404 A–C pairs) and peanut (379 A–B pairs), homoeologs were identified as reciprocal best DIAMOND^41^ hits between subgenome proteomes (homoeogwas prep-homoeologs), restricted to homoeologous chromosome numbers in cotton and peanut and searched across all A–C chromosome pairs in rapeseed; pairs required at least three SNPs per copy. SNPs were mapped to genes within ±2 kb of the gene or mRNA model (±1 kb for peanut; homoeogwas prep-snps), and each mapping file is bound to its genotype file by SHA-256. As a descriptive, genotype-free check of the pairings, we measured local collinearity: genes were ordered by start coordinate, and for each tested edge we counted how many of the ten nearest tested edges along the first copy (five on each side) have partners on the same chromosome within ten positions of its partner in the ordered list of tested partners. Two nulls were used (200 permutations each, seed 20261104): partners permuted within the same chromosome pair, which keeps every partner on its chromosome and the number of pairs per chromosome pair, and partners reassigned freely within the subgenome pair. Per-panel results are given in Supplementary Table 1.

### Phenotypes

Wheat heading date was recorded as days to ear emergence in four field trials (2006, 2010, 2014, 2020)^36^. Line values were the least-squares means of the model value ∼ environment + line, which removes trial effects from lines observed in different trial subsets. For single-locus scans, records more than 6 s.d. from their trial mean were set to missing before adjustment (one record); the interaction analysis and the PVE atlas used the adjusted means without this rule, after inverse-normal transformation. Cotton fibre length and elongation were the best linear unbiased predictors distributed with CottonGVD^37^. Rapeseed flowering time was taken from Wu et al.^40^. Peanut hundred-seed weight and seed length were taken from the PeanutPan study. The other 13 wheat traits of the PVE atlas were single-environment values or least-squares means as above.

### Additive workflow

For each subgenome *s*, a genomic relationship matrix 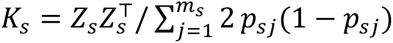was computed from the *m_s_* variants of the subgenome, with *p_s_*_j_ the allele frequency of variant *j* and *Z_s_* the mean-imputed dosages centred by 2*p_s_*_j_^42^, using variants with minor-allele frequency (MAF) ≥ 0.01, and scaled to trace *n*. The model

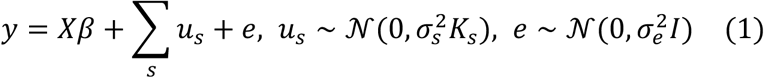

was fitted by restricted maximum likelihood (REML)^43^ with L-BFGS-B from multiple starting points (one deterministic, the others log-uniform; seed 2026; ten by default, five for the PVE atlas), keeping the fit with the highest likelihood. The PVE of subgenome *s* is 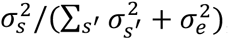; components with PVE below 10⁻³ are flagged as boundary estimates. Uncertainty was quantified by a parametric bootstrap (200 replicates drawn from the fitted model and refitted), giving percentile intervals and the bootstrap frequency with which each subgenome ranks first.

Marker association used the one-degree-of-freedom generalized-least-squares statistic *χ*^2^ = *U*^2^/*I* with variance components fixed at the genome-wide fit; in the leave-one-chromosome-out (LOCO) scan^44,45^ the kernel of the subgenome carrying the tested chromosome is rebuilt without that chromosome’s markers, without refitting the variance components. The genomic inflation factor is median(*χ*^2^)/0.4549^46^.

### Homoeolog-group interaction test

Each group contributes all pairwise edges between its copies (1, 3 or 6 edges for 2, 3 or 4 copies); no higher-order coefficient is fitted. For each copy, SNPs with MAF ≥ 0.01 in the analysed samples were mean-imputed and standardized; for genes with more than 150 such SNPs, separate deterministic subsamples of 150 SNPs were drawn for the burden and for the principal-component features, with seeds derived by SHA-256 from the feature seed, subgenome, gene and feature type. An edge was tested when both copies retained at least three SNPs and its interaction columns were estimable given the covariates and main effects; non-estimable encodings were omitted from the combination, and groups without an estimable edge were excluded from the family. Three encodings of the interaction between copies X and Y were formed: (i) the product of the minor-allele burdens (the mean of standardized SNPs oriented to the minor allele), (ii) the product of the first principal-component scores of the two gene blocks and (iii) up to nine cross-products of up to three unit-variance principal-component scores per copy, tested jointly as fixed-effect interaction columns (a low-rank form of the Hadamard product of the two linear gene kernels).

The phenotype was inverse-normal transformed, Φ^−1^{(*r* − 0.5)/*n*} for rank *r*. The null model 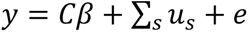 (*C*, intercept) was fitted by REML from three starting points with one relationship matrix per subgenome (standardized SNPs with 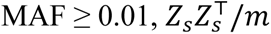 scaled to trace *n*, eigenvalues floored at 10⁻⁶), and the whitening matrix 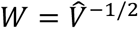 was formed. Each encoding was then tested by a whitened generalized-least-squares nested F-test of its interaction columns, conditional on the covariates and on the main effects of both copies (burden, PC1 or top PCs, respectively), with numerator degrees of freedom equal to the rank added by the interaction columns and denominator degrees of freedom *n* minus the rank of the full design. The three encoding P values of an edge were combined by the Cauchy combination with equal weights^30,47^, 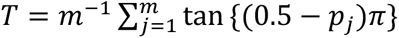 and *P* = 0.5 − arctan(*T*)/*π* for *m* P values, and the edge P values of a group were combined in the same way to give the group P value (group-omniB). Encoding and edge P values are reported to localize the evidence and are not part of the discovery family.

### Experiment-wide error control

All groups of a declared analysis form one family. Null phenotypes

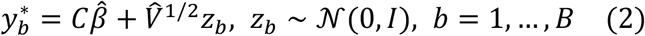

were drawn from the fitted null model, already on the transformed scale, and analysed with the same frozen whitening matrix. For each replicate the minimum P value over the family was recorded; the adjusted P value of group *g* is

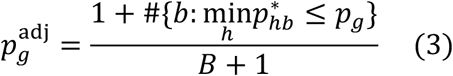

and groups whose P value falls below the ⌊*α*(*B* + 1)⌋-th smallest null minimum are rejected at family-wise level α (single-step minimum-P)^31^. Replicates with a non-finite statistic contribute a minimum of zero, which is conservative. We used B = 2,000 and α = 0.05 throughout.

In the smooth_pc4 null, the bootstrap residuals are rescaled by a smooth variance function of population structure. The squared whitened residuals of the null fit were regressed on an intercept and the top four eigenvectors of the mean subgenome relationship matrix by a gamma generalized linear model with log link (iteratively reweighted least squares, at most 50 iterations); the fitted variances *w*, normalized to mean one, define

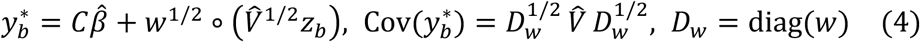

The weights therefore rescale the whole fitted covariance, including its relationship components, sample by sample; because they are estimated from whitened residuals, they are a working approximation to this structure rather than a likelihood estimate, and their adequacy was assessed by simulation under two forms of heteroscedasticity (below). The observed statistics are unchanged. The homoscedastic null (*w* = 1) is available as a sensitivity analysis. smooth_pc4 was selected after the unmodified bootstrap and two wild bootstraps (Rademacher and Mammen weights) failed internally frozen criteria in a pilot, and was then validated on fresh phenotypes with criteria frozen before analysis.

### Calibration and power experiments

#### Families and generators

Calibration and conditional power used, per panel, an 80-group family selected by stratifying on chromosome and marker-count quartile with a hash-based order that used no phenotype; larger families of 500 and 2,000 groups nest the 80 groups. Simulated signals were pairwise copy-by-copy products, standardized: minor-allele burden product, PC1 product, multidimensional kernel (sum of products of the first ≤ 3 PCs), single SNP pair, alternating-sign SNP sums and, for wheat triads, burden products on two edges. For a target PVE *q*, the signal *s* was standardized, the background was centred, projected orthogonally to *s* and standardized (*e*_⟂_), and 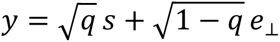, so that PVE is the exact finite-sample share of the untransformed phenotype variance. The additive-only control adds per-copy PC1 main effects without interaction; the mispaired control is the burden of one copy times a gene of another group.

#### End-to-end calibration

In wheat and cotton, 1,250 Gaussian null phenotypes with the panel’s kinship covariance and 500 each with an added additive or PC1 main effect were analysed by the complete pipeline (per-phenotype REML, B = 2,000) with the homoscedastic null; rapeseed was analysed in the same way with smooth_pc4. Failures counted as rejections, and the prespecified criterion was a Wilson upper bound ≤ 0.075.

#### Variance-model validation

In wheat, cotton and rapeseed, the homoscedastic and smooth_pc4 bootstrap arms were applied to the same fresh phenotypes, generated independently of the development pilot, under Gaussian, Student-t (5 d.f., unit variance), 1% contamination at ±6 s.d., PC1-aligned heteroscedastic (the whole kinship-preserving draw scaled by a variance multiplier 1 + 3 minmax(PC1), matching the form assumed by smooth_pc4) and non-PC heteroscedastic nulls (2,000 phenotypes per layer, 500 for the last). The frozen criteria were a Wilson upper bound ≤ 0.075 for smooth_pc4 in each core layer and a paired power loss ≤ 0.05 in 12 positive cells (observed maximum 0.022).

#### Threshold transfer

For each panel, statistic and null layer — Gaussian and additive-only (core), Student-t, PC1-heteroscedastic and contaminated (stress), and an omitted-kernel diagnostic — the rejection threshold (100th smallest family minimum P among 2,000 calibration phenotypes) was applied to 2,000 held-out phenotypes, using the conditional test with a fixed whitening matrix and no bootstrap (B = 0). The same banks provided the 80-group power grid (500 replicates per cell; 1 or 4 causal groups; PVE 2%, 5%, 10%) and the comparisons of group-omniB with each of the three single encodings in 96 positive cells (288 comparisons), each statistic at its own threshold.

#### Whole-family calibration and power

Null phenotypes *y* = *u* + *e* with *u* ∼ N(0,0.30*K*) and *e* ∼ N(0,0.70*D*), where *K* is the mean subgenome relationship matrix (mean diagonal one) and *D* = diag(*d_i_*/*d̄*) with *d_i_* = exp{0.4 *z*(PC1*_i_*) + 0.4 *z*(PC2*_i_*)} for the first two eigenvectors of *K* and *d̄* the mean of *d_i_* (heteroscedasticity in the residual component only, a form different from the one assumed by smooth_pc4), were analysed over each complete family with the delivered pipeline (INT, per-phenotype REML, smooth_pc4, B = 2,000; 640 phenotypes per panel). The prespecified criterion was a one-sided 95% Clopper–Pearson upper bound ≤ 0.075. For power, in wheat, cotton and rapeseed (peanut entered the whole-family calibration but not the power experiment), one interaction signal was added on this background to a unit drawn uniformly among units whose copies each retained at least three SNPs (all units of each family): six architectures in wheat and five in each two-copy panel, at PVE 2%, 5% and 10% (48 cells × 200 phenotypes = 9,600 analyses). The background was projected orthogonally to the signal so that PVE is the exact share of the untransformed phenotype variance before inverse-normal transformation. The signal used all gated SNPs of a gene, whereas the engine scores at most 150; genes with more than 150 mapped SNPs were 4.5% of tested gene copies in wheat and at most 0.4% in the other panels. Non-causal rejections were classified, in order, as on a chromosome carrying any copy of the causal unit, on another chromosome with the same number, or elsewhere. Family scaling reanalysed the same 500 Gaussian and PC1-heteroscedastic phenotypes per panel in wheat, cotton and rapeseed at 500 and 2,000 groups (smooth_pc4).

#### SNP-by-SNP comparator

In the 80-group families of wheat, cotton and rapeseed, regenie 4.1.2^48^ was run per phenotype with whole-genome ridge regression and leave-one-chromosome-out predictions (step 1) and a test of the product of each within-group cross-copy SNP pair with both main effects (step 2, –interaction-snp –rare-mac 0 –no-robust); the group score was the minimum pair P value. Group-omniB used the observed group P value (fixed whitening, B = 0). Both methods were thresholded at the 100th smallest family minimum of 2,000 Gaussian calibration phenotypes and evaluated on 2,000 held-out phenotypes, on additive-only and mispaired controls and on the same 320 phenotypes per positive cell; paired power differences have Newcombe 95% intervals. regenie excludes samples missing at a tested SNP, whereas HomoeoGWAS mean-imputes.

#### Additive-effect conditions

The analyses with additive effects and no interaction differ in the additive effect (a genome-wide genotype or PC1 main effect, per-copy PC1 effects of one control group, or a planted variant outside the tested blocks), in the phenotype scale, in whether whitening is fixed or refitted per phenotype, in the threshold source and in whether the bootstrap is used; Supplementary Table 15 lists the conditions and rates side by side.

#### Location of control rejections

In an exploratory analysis of the stored conditional-grid results, each rejected group under the additive-only and mispaired controls was classified as the control group, the group of the mispaired partner gene, another group on the same homoeologous chromosome number, or elsewhere (one control group, 10% PVE, group-omniB).

#### Refit diagnostic

In an exploratory analysis in wheat and cotton, stored phenotypes were rescored with the fixed whitener on the untransformed or transformed response and with per-phenotype REML on the transformed response (200 phenotypes per control, 500 Gaussian; each variant with its own 1,000-phenotype calibration).

All confirmatory simulation designs were prespecified: code and parameters were frozen with hashes in version control before the corresponding results were generated, and amendments affected execution only. These protocols were not publicly registered. Seeds were derived from fixed roots by SHA-256 or numpy SeedSequence.

### Application analyses

Each application trait was analysed with one family and one bootstrap-minP correction (group-omniB; configured at the edge level in rapeseed and at the group level otherwise, which coincide for two-copy groups; burden cap 150, minimum three SNPs per copy, MAF ≥ 0.01, three PCs; smooth_pc4; B = 2,000; bootstrap and feature seed 2026).

#### Known loci

Recovery was assessed in fixed windows around loci reported for the same panels or around cloned genes: cotton fibre-length loci of Peng et al.^37^ (±250 kb), 29 heading-date loci of Cheng et al.^36^ (±5 Mb; expectation and P value from 1,000 joint per-chromosome circular shifts), seven rapeseed flowering loci (±100 kb; Poisson-binomial expectation for random windows) and the peanut seed genes AhARF2-2 and AhCKX6 (±100 kb). We report genome-wide Bonferroni significance (wheat 6.0 × 10⁻¹⁰, cotton 4.52 × 10⁻⁸, rapeseed 4.95 × 10⁻⁸, peanut 8.51 × 10⁻⁸) and, for rapeseed and peanut, P < 10⁻⁵. Wheat was also scanned for the 2006 trial alone, and peanut counts were compared with GEMMA 0.98.5^18^ on the same data.

#### PVE atlas

Fourteen inverse-normal-transformed wheat traits were fitted with A, B and D kernels built from LD-pruned SNPs (plink2 –indep-pairwise 1000kb 1 0.2; MAF 0.01; about 90,000–95,000 SNPs per subgenome), five starting values for the observed fit and 200 bootstrap replicates with one start each; four candidate traits were re-audited with five starts per replicate. The prespecified gate required convergence with at least 80% successful bootstrap refits, a maximum PVE change ≤ 0.10 across MAF thresholds (0.01, 0.05) and transformations, the same top subgenome in models with and without a homoeolog kernel (the elementwise product *K_A_* ∘ *K_B_* ∘ *K_D_*), a top-rank frequency ≥ 0.80 in both models and bootstrap contrasts with the other subgenomes excluding zero. The D share of starch genetic variance was compared with 1,000 random groupings of the 21 chromosomes into three sets of seven.

#### Robustness

Accession deletion used 20 deterministic folds and, for wheat, leave-one-environment reanalyses with the environment-adjusted mean refitted from the remaining trials; the interaction features were held fixed and nominal P values are reported. In the additive-leakage control, a variant with MAF ≥ 0.05 within ±5 Mb of a detected unit, outside every tested gene block and maximally correlated with the unit’s burden and PC1 edge products, was added with PVE 0, 10% or 20% to the whole-family null generator (640 phenotypes per cell; criterion as for the whole-family null); a detected unit was to be withdrawn if rejected in at least 7 of 640 replicates. Regional conditioning reran each family with the top five genotype PCs per copy within ±1 Mb (sensitivity: ten PCs within ±5 Mb) as covariates; a unit was retained when the Clopper–Pearson upper bound of its adjusted P was below 0.05 in both windows and not robust when the lower bound exceeded 0.05 in either. Three-copy conditional models of wheat heading date (stable A × B × D after all lower-order terms on the line mean; environment × A × B × D after all lower-order environment interactions on 3,099 line × environment observations with a line-block Rademacher wild bootstrap; α = 0.025 each) were taken from their frozen analyses; the second was re-executed from its frozen launcher. Candidate genes were annotated by orthology and protein domains (InterPro^49^), and expression was taken from the wheat developmental atlas^8^.

#### Unit checks

For each significant unit we recorded the strongest additive SNP P value within each copy (gene ± 20 kb) and within ±1 Mb, and the absolute correlation between each copy’s PC1 feature and the top five genotype principal components, expressed as a percentile among all units of the family and among the 50 units nearest in gated SNP count and mean MAF (Mahalanobis distance).

### Homoeolog coupling

Per-copy PC1 features were replayed from the formal interaction runs (reproducing the formal P values to 10⁻¹¹) and residualized on ten genotype principal components computed from 20,000 random gated SNPs. Homoeolog coupling *r_P_* of an edge *e* is the absolute Pearson correlation |*r_e_*| of the two residualized features. With *z_X_* and *z*_F_ the centred, unit-norm feature vectors over all *n* accessions of the panel, the per-accession contribution *s*(*e*, *i*) = sign(*r_e_*) *n z_X_*_,*i*_*z*_F,*i*_ averages to *r_P_*. In a kinship-whitened sensitivity analysis the features were premultiplied by *V*^−1/2^ of a genome-wide relationship model before correlation^50^. The null re-paired each copy with a copy of another unit on a different chromosome number (1,000 re-pairings). Reliability was estimated from 20 splits of the panel into halves along k-means clusters (k = 40) of the principal components, as the median covariate-conditioned Spearman correlation between half-panel edge values; its lower bound (5th percentile of 1,000 5-Mb block bootstraps) was projected to full-panel scale by the Spearman–Brown formula and compared with the minimum reliability needed for 80% power to detect ρ = 0.13 in an external edge-level test.

Rapeseed homoeologous exchange (HE) segments for 260 accessions were taken from Wang et al.^29^. A copies were linked to the study’s anchor genes through one-to-one BnPIR orthogroups^51^, and the HE frequency of an edge is the number of accessions with a segment covering the anchor’s C-gene midpoint; exchange direction (A replacing C or C replacing A) was taken from each segment’s annotation in that study. Associations were tested by Spearman correlation against a within-chromosome circular-shift null: edges were ordered by A-copy position, the HE frequencies were circularly shifted within each A chromosome (2,000 shifts), and the one-sided P value is (1 + number of shifted ρ ≥ observed)/(2,000 + 1), so that 5 × 10⁻⁴ is its floor; covariate-adjusted partial correlations included SNP count, MAF and heterozygosity. In rapeseed, filtering heterozygous calls removes most of the genome-wide coupling excess over re-pairing, so the association with HE frequency was also tested after adjustment for per-edge heterozygosity. For the 123 accessions shared with that study (identical names and sequencing runs), the carrier statistic is the difference in mean contribution between carrier and non-carrier accession–edge pairs, with a null that reassigns whole HE profiles among accessions of the same ecotype (1,000 permutations). In a post hoc analysis, edges were stratified by coupling strength (*r_P_* < 0.1, 0.1 ≤ *r_P_* < 0.3 and *r_P_* ≥ 0.3), and the carrier difference was recomputed within each stratum against 300 permutations of the same null, with P = (1 + number of permuted differences ≥ observed)/301. Intervals for mean coupling by HE-frequency bin are percentile intervals of 1,000 bootstrap resamples of edges within each bin; intervals for the mean contribution of carriers and non-carriers are percentile intervals of 1,000 resamples of whole accessions, each resampled accession bringing all of its accession–edge pairs. External wheat and cotton tests used introgression and copy-number frequencies^36,52^, pan-genome gene classes^53^, homoeolog expression co-variation^54,55^ and *G. barbadense* introgression and presence–absence variation^56^, each against spatial circular-shift nulls. Cotton within-group tests used the 1,170 accessions assigned to cultivar groups Cul1 (*n* = 449) and Cul2 (*n* = 721) by Xu et al.^57^: coupling was recomputed within each group with ten group-specific principal components, its excess over 1,000 density-matched re-pairings (partner drawn from the same SNP-count quintile) was combined across groups with inverse-variance weights, and the one-sided P value used the same permutation index in both groups. Each primary test and statistic was frozen in version control before it was computed; post hoc analyses are labelled as such. The reliability threshold was set after the genome-wide coupling summaries were available, before any split-half statistic was computed, and the reliability assessment is therefore exploratory.

### Computational benchmark

Timing used two nodes, each with two 64-core AMD EPYC 7742 processors (1,007 GB and 503 GB memory; single-locus scans and mvMAPIT on the first, the other interaction analyses on the second), one cell at a time, with /usr/bin/time -v and three repeats (median and range; the first repeat with a cold cache; four repeats for mvMAPIT; 24-h cap). Repeats overlapping other heavy jobs are flagged and reported. Single-locus scans of wheat heading date (827 accessions; 0.1, 1, 10 and 83.3 million nested random markers; 64 threads) compared HomoeoGWAS (A, B and D kernels with LOCO; streaming in chunks of 200,000 markers), GEMMA 0.98.5 (one kinship; -gk 1, -lmm 1)^18^, REGENIE 4.1.2 (step 1 block size 1,000)^48^ and PLINK 2.0.0-a.6.9 (–glm, no kinship)^58^. Panels of 2,500 and 5,000 genotypes were simulated for timing by copying chromosome segments (mean 20 Mb) from real lines, with a phenotype of heritability 0.5. Interaction analyses used 128 HomoeoGWAS workers with one native thread each; PLINK 1.9.0-b.8 –epistasis (all SNP pairs of the selected groups), REGENIE within-group SNP-pair tests and mvMAPIT 2.0.4^59,60^ used 64 threads; the PLINK 1.9 time includes writing about 0.95 TB of output to network storage. B = 19 is the smallest bootstrap size the engine accepts and was used to estimate the cost of calibration.

### AI-assistant evaluation

Thirty frozen tasks (10 routine analyses, 8 input traps, 7 invalid requests and 5 interpretation questions; data cut from the real panels) were given to Claude Code CLI 2.1.296 with model claude-sonnet-5-5 and Codex CLI 0.162.1 with model gpt-6-sol in isolated sandboxes with (A) the HomoeoGWAS skill and tool server, (B) the command-line interface and documentation, or (C) no tools (invalid-request and interpretation tasks only), three repeats each (432 runs). Routine tasks were scored deterministically against expert reference outputs; other tasks combined deterministic checks with an isolated claude-sonnet-5-5 judge, cross-checked on 311 runs by a gpt-6-sol judge run through Codex CLI (agreement 0.913, κ = 0.818). Task definitions, prompts and scoring rubrics are provided in the reproducibility archive. A before–after comparison of the software without and with the added input guards used three input-trap tasks (36 runs per release) and two valid control tasks (24 runs per release) across both agents and arms A and B (120 runs).

### Software and reproducibility

HomoeoGWAS is written in Python and released under the MIT licence (GitHub, PyPI and documentation site). A preflight refuses to run on missing genotype files, a SNP-to-gene map whose genotype hash does not match, duplicated sample or gene identifiers, mismatched sample order, fewer than ten overlapping samples or an undeclared marker encoding. Every formal run records a pre-run manifest with the configuration hash, the git commit and a clean-tree check; checkpoint manifests bind the family, phenotype, samples, relationship matrices, features and null fit by SHA-256. Parallel interaction analyses use forked workers with one native linear-algebra thread each, and experiment-wide decisions were identical across worker counts in a wheat check. The exact software version used for each analysis is recorded in the Zenodo archive. Configurations, frozen analysis protocols, figure source data and per-replicate outputs will be deposited in a versioned Zenodo archive.

## Data availability

The genotype and phenotype data analysed in this study are publicly available. Genotypes and heading-date records of the Watkins wheat collection are available from the European Nucleotide Archive under accession PRJEB71453^36^. Genotypes and fibre-quality phenotypes of the cotton panel are available from CottonGVD (https://db.cngb.org/cottonGVD/)^37^. Rapeseed genotypes are available from BnaGVD (https://rapeseed.biocloud.net/)^38^, and flowering-time records were taken from Wu et al.^40^. Peanut genotypes are available from PeanutPan (https://cgm.sjtu.edu.cn/PeanutPan/), and seed-trait phenotypes were taken from Zhao et al.^39^. Homoeologous-exchange segments of *B. napus* were taken from Wang et al.^29^; the additional wheat and cotton datasets used for external comparisons are cited in Methods. The source data underlying the figures, together with simulation configurations, analysis protocols and per-replicate outputs, will be deposited in a versioned Zenodo archive.

## Code availability

HomoeoGWAS is open-source software under the MIT licence. The current release (v2.2.0; Python ≥ 3.10) is available from GitHub (https://github.com/Shipeng-Yang/HomoeoGWAS, release v2.2.0) and PyPI (https://pypi.org/project/homoeogwas/2.2.0/), with documentation at https://shipeng-yang.github.io/HomoeoGWAS/. All releases are archived at Zenodo (https://doi.org/10.5281/zenodo.20668367). Scripts that reproduce the simulations, application analyses, AI-assistant evaluation and every figure will be made available at https://github.com/Shipeng-Yang/HomoeoGWAS-reproducibility and archived in the same Zenodo archive.

## Acknowledgements

We thank Claude Code (Anthropic) for assistance with software development, code and manuscript review, and language editing; the authors checked all outputs and take full responsibility for the content.

## Author contributions

S.Y. conceived the study, developed HomoeoGWAS, performed the analyses and wrote the manuscript. Q.Z. co-supervised the study and revised the manuscript. C.S., Y.W., Z.S. and C.Y. contributed to data collection and curation. W.L., H.Z., L.W. and X.S. contributed to the interpretation of the results. All authors read and approved the final manuscript.

## Supplementary Information

### HomoeoGWAS enables experiment-wide homoeolog-group interaction mapping in allopolyploid crops

Contents: Supplementary Figs. 1–9; Supplementary Tables 1–15 (provided as Supplementary_Tables_1-15.xlsx); Supplementary Data 1 (SupplementaryData1.xlsx); Supplementary References.

## Supplementary Figures

**Supplementary Fig. 1.**
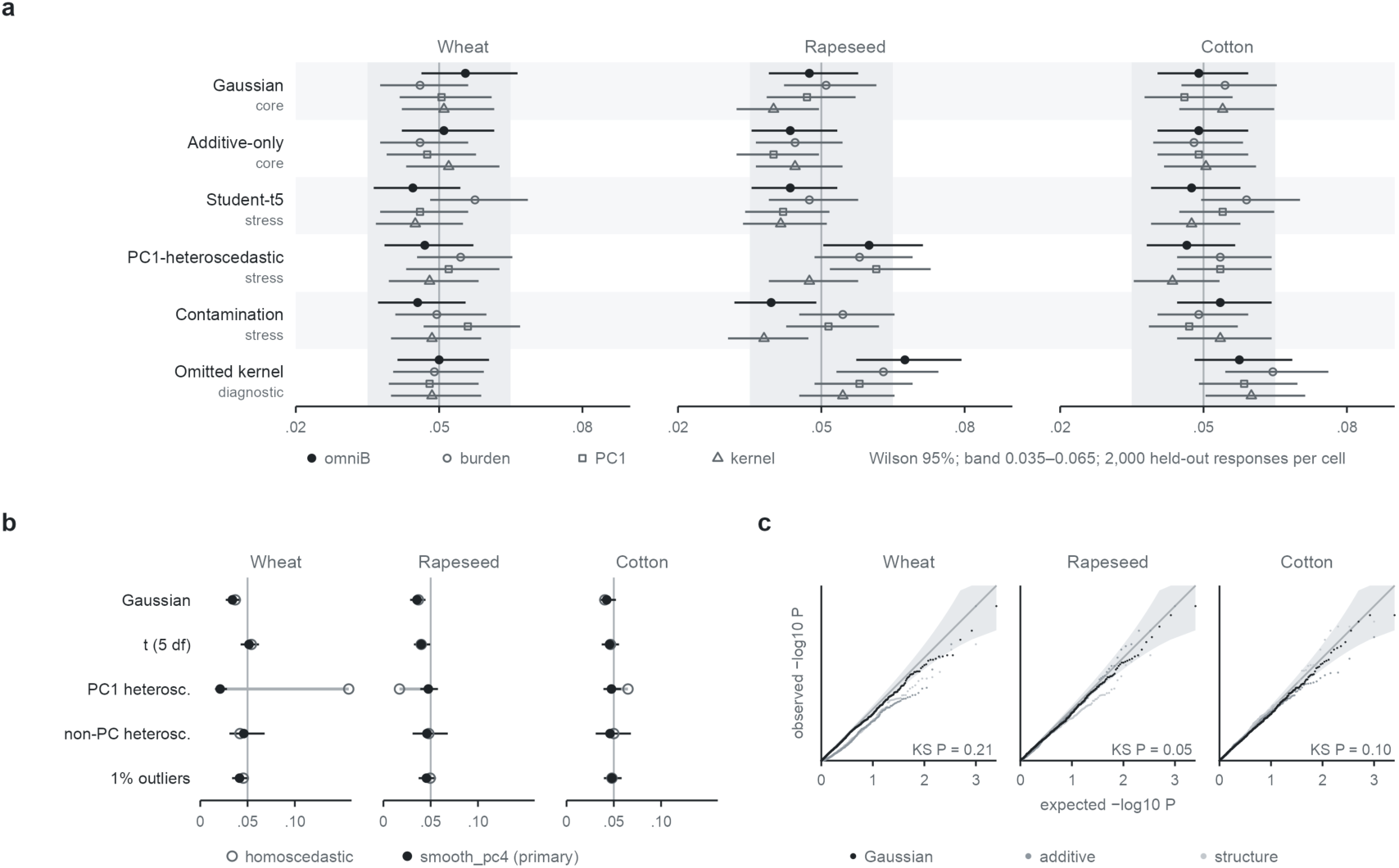
Calibration of the interaction test in wheat, rapeseed and cotton. **a**, Threshold transfer: held-out FWER of the conditional test (fixed whitening, B = 0) for group-omniB (filled circles) and the burden (open circles), PC1 (squares) and kernel (triangles) encodings under six null layers; thresholds from 2,000 calibration phenotypes per layer, 2,000 held-out phenotypes per cell. **b**, Variance model: FWER of bootstrap-minP (B = 2,000) under five noise models; open, homoscedastic null; filled, smooth_pc4. **c**, Q–Q plots of the global P under Gaussian (black), additive (dark grey) and structure (light grey) nulls; shading, pointwise 95% interval; KS P, Kolmogorov–Smirnov test of the Gaussian-null global P against uniform. Bars, Wilson 95% intervals; vertical line, 0.05; grey band, 0.035–0.065.

**Supplementary Fig. 2.**
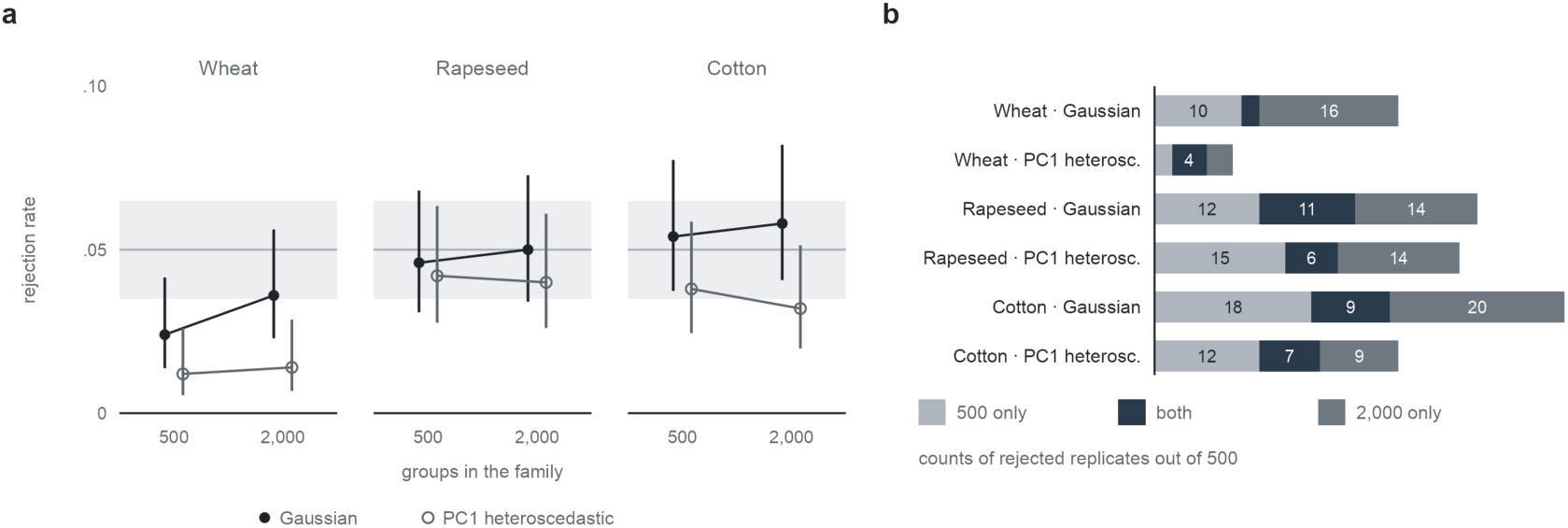
Family-size scaling under the global null. **a**, Any-group rejection at 500 and 2,000 groups (smooth_pc4, B = 2,000) under Gaussian (filled) and PC1-heteroscedastic (open) nulls, the same 500 phenotypes per layer at both sizes; bars, Wilson 95% intervals; grey band, 0.035–0.065. **b**, Phenotypes rejected at 500 groups only, at both sizes or at 2,000 groups only.

**Supplementary Fig. 3.**
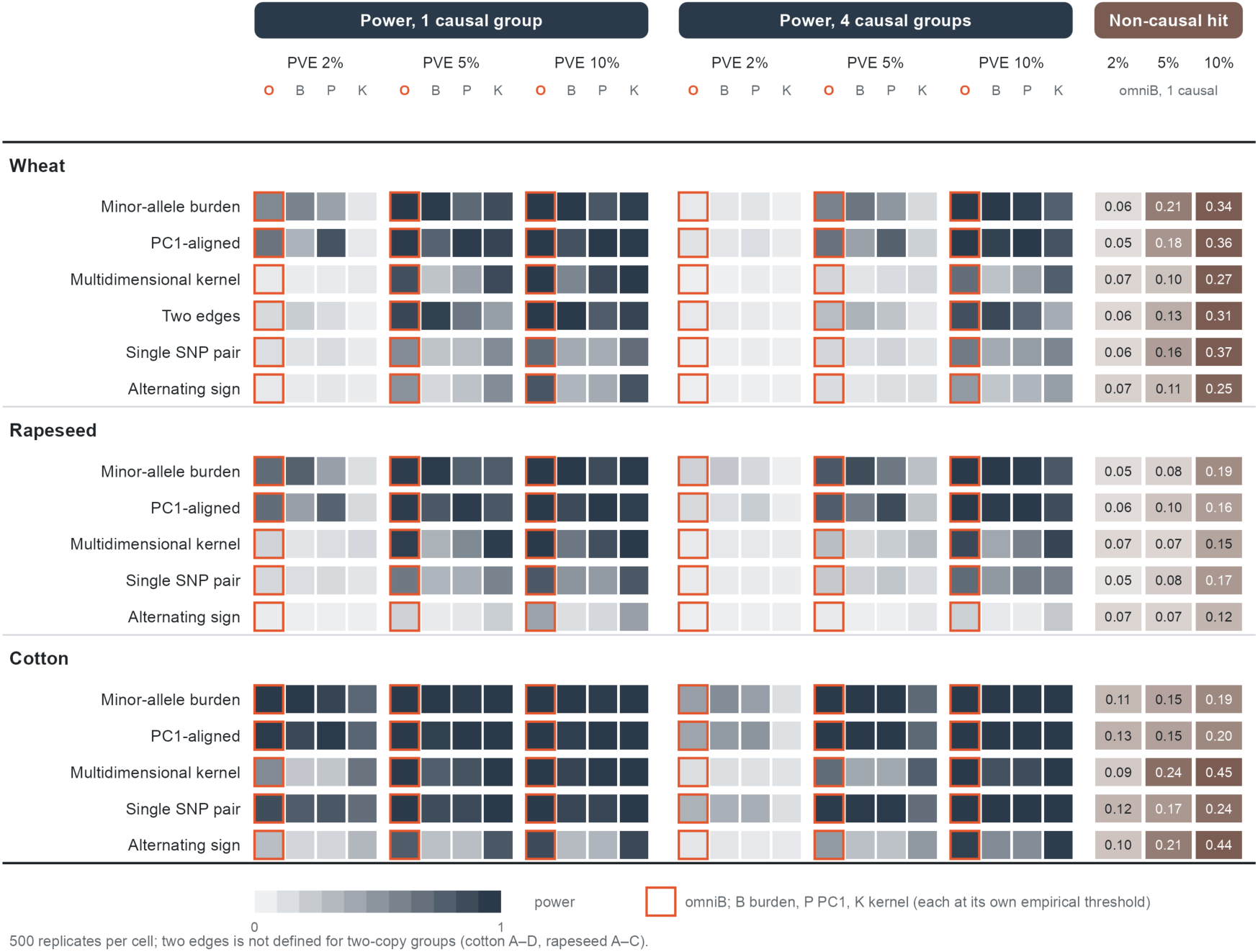
80-group conditional power grid. Power (shading) of group-omniB (O, orange outline) and the burden (B), PC1 (P) and kernel (K) encodings for each architecture, PVE and number of causal groups; 500 replicates per cell; each statistic at its own empirical threshold. Right, fraction of group-omniB replicates with one causal group in which a non-causal group was rejected.

**Supplementary Fig. 4.**
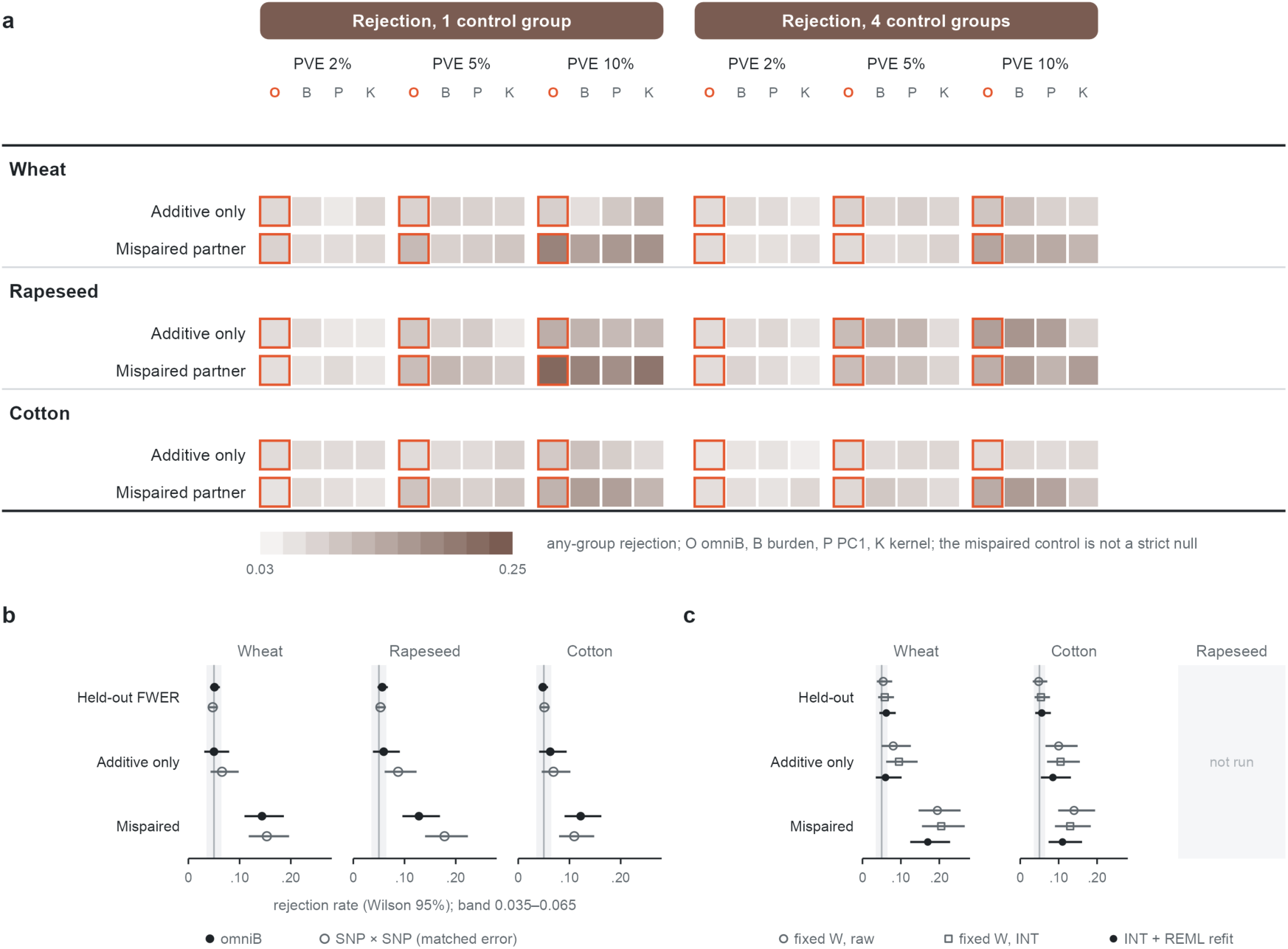
Control architectures, comparator controls and refit diagnostic. **a**, Any-group rejection (shading) under the additive-only and mispaired-partner controls for group-omniB (O) and the burden (B), PC1 (P) and kernel (K) encodings; 500 replicates per cell. **b**, Held-out FWER (n = 2,000) and additive-only and mispaired controls (PVE 10%, n = 320) for group-omniB (filled) and the SNP × SNP scan (open) at matched error. **c**, Group-omniB rejection in wheat and cotton with a fixed whitening matrix on the untransformed (open circles) or transformed phenotype (squares) and with per-phenotype REML on the transformed phenotype (filled); held-out n = 500, controls n = 200. Bars, Wilson 95% intervals; grey band, 0.035–0.065.

**Supplementary Fig. 5.**
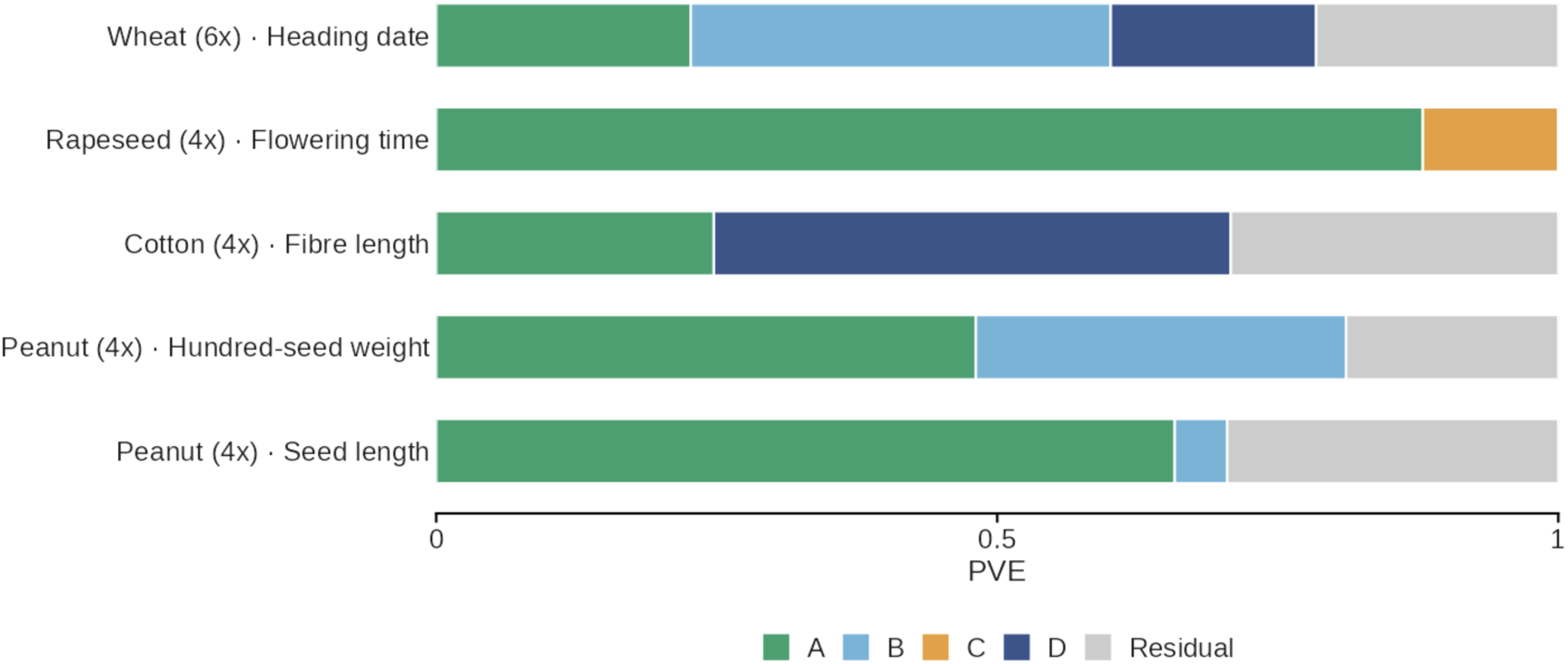
Subgenome partition of the application traits. REML PVE of each subgenome kernel (colours) and the residual (grey) for the scans of Fig. 3a.

**Supplementary Fig. 6.**
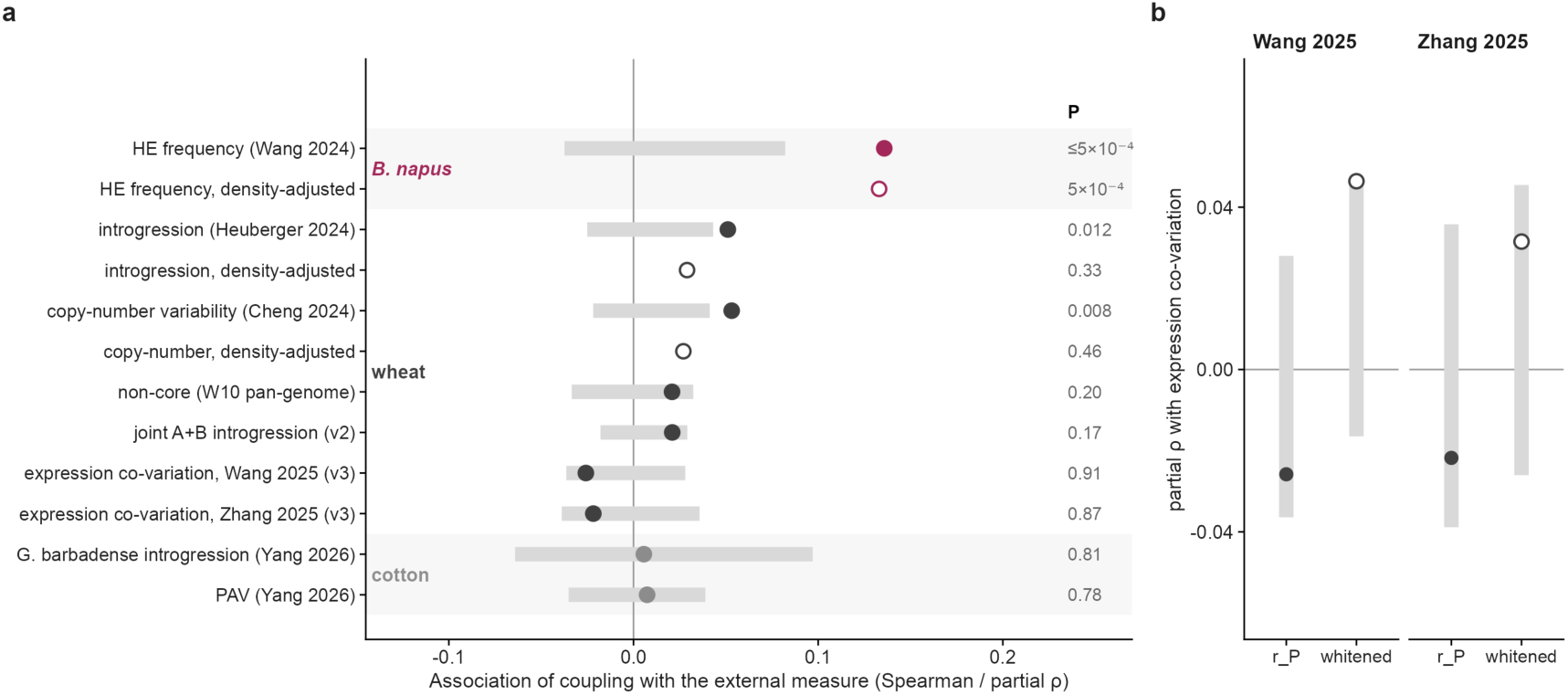
External tests of homoeolog coupling. **a**, Association of edge coupling *r_P_* with external measures^1–7^ (Spearman or partial ρ; rank-biserial for the non-core gene class). Filled, prespecified primary or replication test; open, post hoc density-adjusted reanalysis; colour, species; grey bars, 95% range of the within-chromosome circular-shift null; right, permutation P. **b**, Wheat: partial ρ with expression co-variation for *r_P_* (filled) and kinship-whitened coupling (open); grey bars, 95% null range.

**Supplementary Fig. 7.**
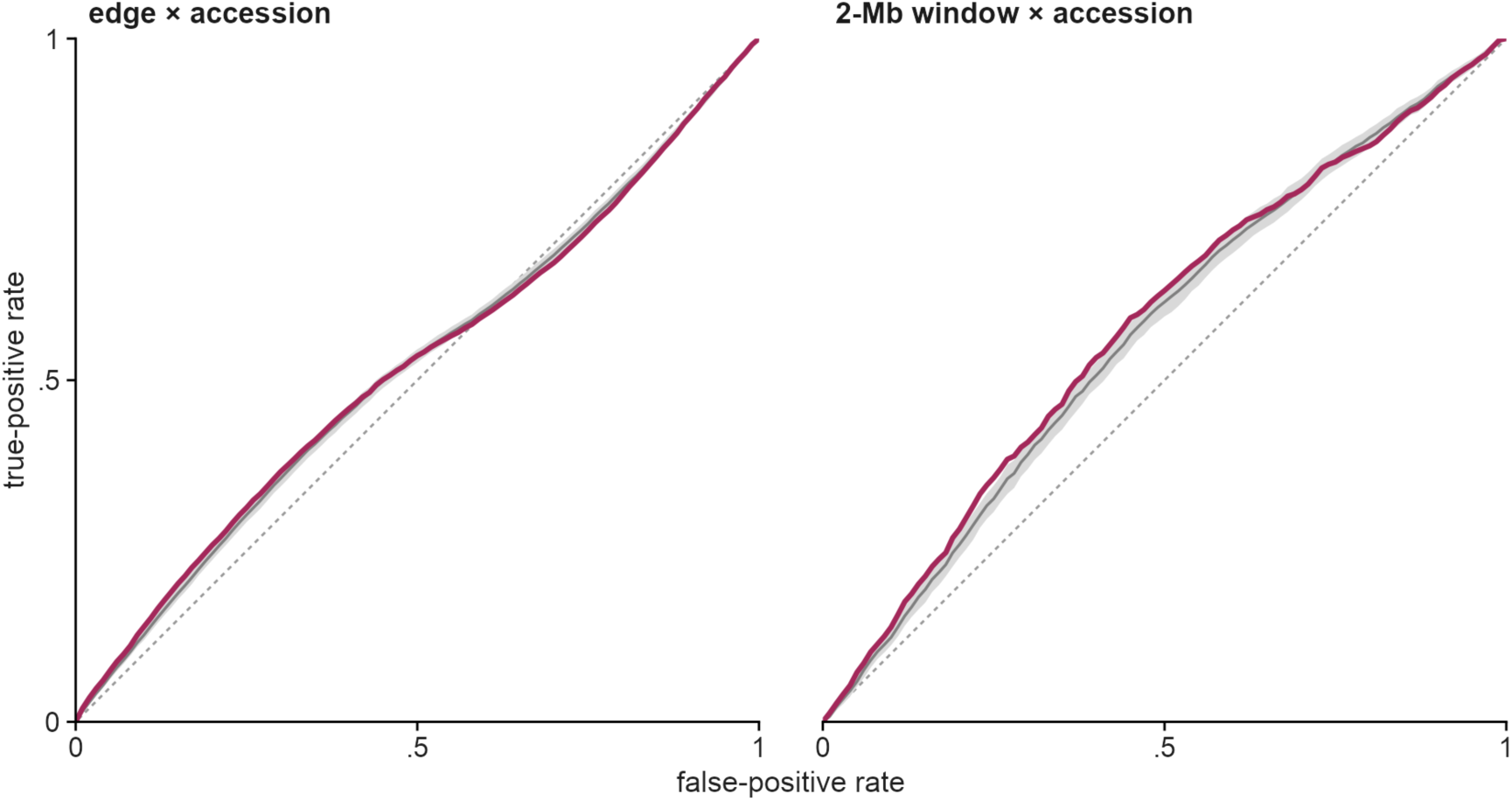
Per-accession HE detection by coupling contribution. ROC curves of the per-accession contribution for HE carrier status in the 123 shared *B. napus* accessions^1^, per edge × accession (left) and per 2-Mb window × accession (right). Magenta, observed; grey line and band, mean and 95% range of 200 within-ecotype reassignments; dotted line, chance.

**Supplementary Fig. 8.**
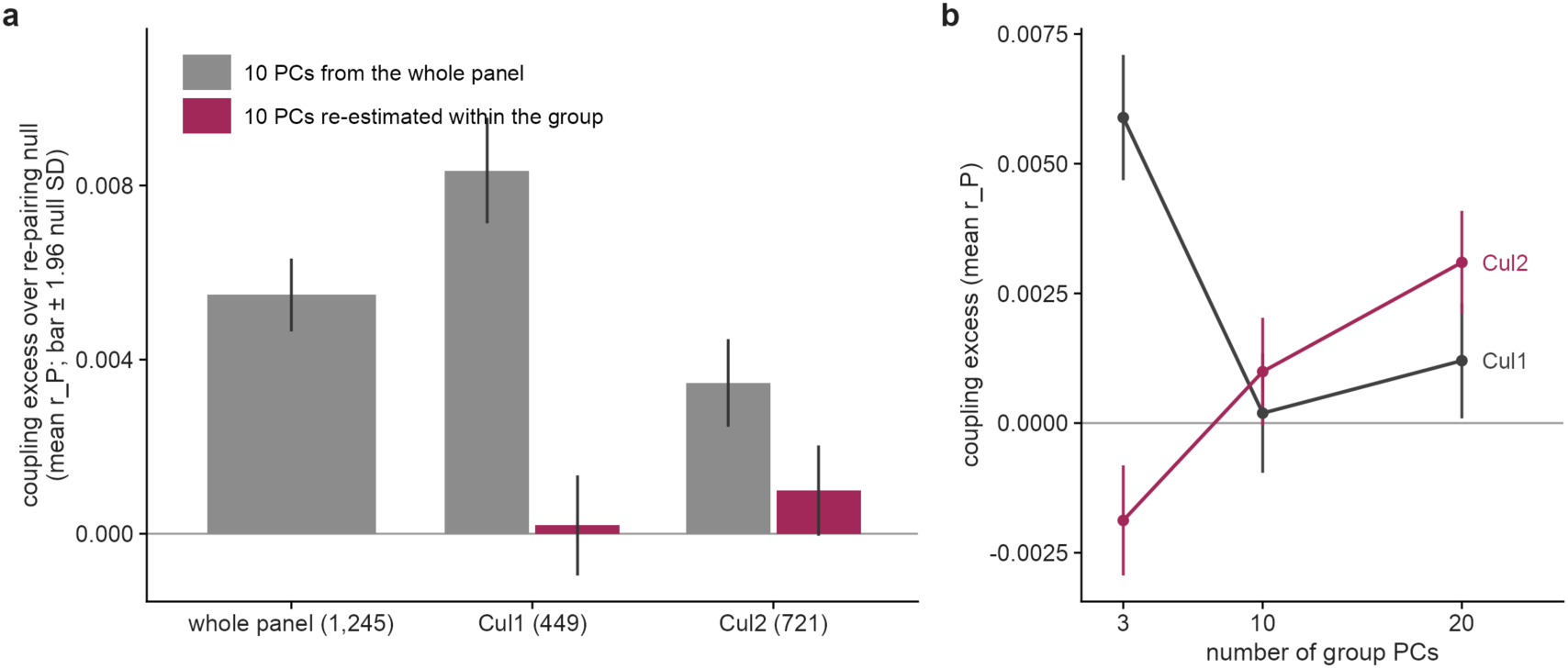
Cotton coupling and population structure (post hoc). **a**, Coupling excess over a density-matched re-pairing null in the whole panel and in cultivar groups Cul1 and Cul2^8^, with ten principal components from the whole panel (grey) or re-estimated within the group (magenta). **b**, Excess with 3, 10 and 20 within-group principal components. Bars, ±1.96 s.d. of 200 re-pairings.

**Supplementary Fig. 9.**
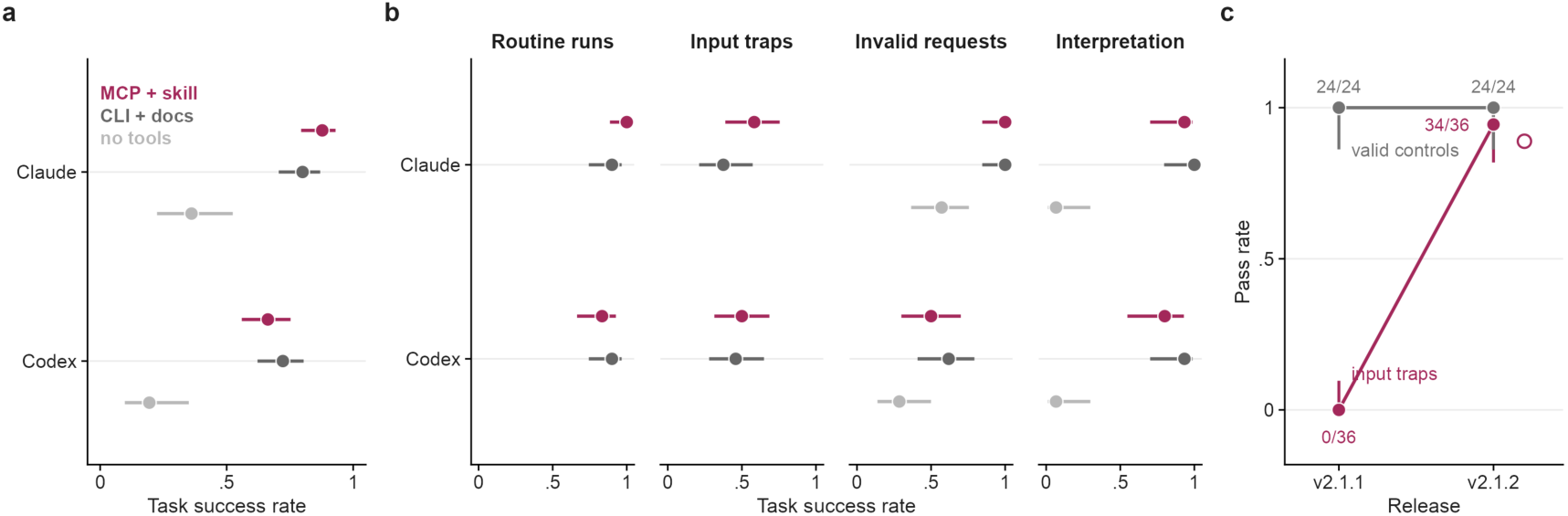
AI-assistant interface evaluation. **a**, Task success rate of Claude Code and Codex CLI on 30 frozen tasks with the HomoeoGWAS skill and MCP tools (magenta), the command-line interface and documentation (dark grey) or no tools (light grey); 432 runs. **b**, Success rate by task category. **c**, Pass rate of input-trap (magenta, n = 36) and valid control tasks (grey, n = 24) without (v2.1.1) and with (v2.1.2) the added input guards; open circle, strict scoring. Bars in a and b, Wilson 95% intervals.

## Supplementary Tables

Supplementary Tables 1–15 are provided as one workbook (Supplementary_Tables_1-15.xlsx), one sheet per table.

**Supplementary Table 1 | Local collinearity of the tested homoeolog families and of the nine detected units.** Support, number of the 10 nearest family edges along the first copy whose partners lie on the same chromosome within ±10 family positions of the edge’s partner. Nulls (200 permutations each, seed 20261104): free, partners reassigned among all edges of the same subgenome pair; constrained, partners permuted within the same chromosome pair. Bottom, the nine detected units.

**Supplementary Table 2 | Whole-family global-null calibration of the four interaction families.** Null phenotypes y = u + e, u ∼ N(0, 0.30K), e ∼ N(0, 0.70D), D = diag(d_i / mean d), d_i = exp{0.4 z(PC1_i) + 0.4 z(PC2_i)}; delivered pipeline (INT, per-phenotype REML, smooth_pc4, B = 2,000, α = 0.05); prespecified criterion: one-sided 95% Clopper–Pearson upper bound ≤ 0.075.

**Supplementary Table 3 | Recovery of known loci by the subgenome-stratified scans.** a, cotton fibre length; b, rapeseed flowering time; c, wheat heading date; d, peanut seed traits. Windows and expectations as in Methods.

**Supplementary Table 4 | Subgenome PVE atlas of 14 wheat traits.** REML with A, B and D kernels; 200 parametric bootstrap fits per trait; screen, screening fits; multistart, audited fits; gate_passed, prespecified stability gate (Methods).

**Supplementary Table 5 | Homoscedastic versus smooth_pc4 bootstrap null for every interaction analysis.** B = 2,000, seed 2026, α = 0.05, one bootstrap-minP family per analysis; smooth_pc4 is the primary null. Observed P values are identical under the two nulls; only the threshold changes. Cells: global P / threshold / rejected groups.

**Supplementary Table 6 | Checks of the nine detected units against nearby additive signal and population structure, and candidate genes.** a, unit summary; b, percentiles among all units (pct_B) and among 50 matched units (pct_M); c, allele-frequency differentiation; d, class tests; e, candidate genes of the four wheat triads (smooth_pc4).

**Supplementary Table 7 | Planted additive-leakage control and regional conditioning.** Planted additive variant at 0, 10 and 20% PVE (cells C0, C10, C20; 640 phenotypes each) and regional conditioning of the nine detected units (±1 Mb with five PCs per copy; ±5 Mb with ten PCs per copy).

**Supplementary Table 8 | Accession- and environment-deletion reanalyses.** Nine detected units with the formal features fixed: 20 deterministic accession folds (all panels) and leave-one-environment reanalyses (wheat); nominal P values.

**Supplementary Table 9 | Wheat triads significant only under the homoscedastic null (sensitivity tier).** Triads significant under the homoscedastic null; the six marked “homoscedastic only” form the sensitivity tier.

**Supplementary Table 10 | Wheat matched edge family and per-direction analysis.** Rejected edges of the wheat matched edge family (smooth_pc4) and analyses of each subgenome direction (AB, AD, BD) with the environment-adjusted phenotype.

**Supplementary Table 11 | Wheat three-copy conditional models.** Prespecified three-copy conditional models of wheat heading date (days to ear emergence) over 2,143 triads; α = 0.025 each.

**Supplementary Table 12 | External tests of homoeolog coupling.** Associations of homoeolog coupling with external measures; prespecified primary or replication tests and post hoc analyses are labelled. Null mean and s.d. of the within-chromosome circular-shift null where available.

**Supplementary Table 13 | Computational benchmark: timing, memory, thread budgets and tool versions.** Median and range of repeats; peak resident memory; declared threads; interference flags. Tools: HomoeoGWAS, GEMMA 0.98.5, REGENIE 4.1.2, PLINK 2.0.0-a.6.9, PLINK 1.9.0-b.8, mvMAPIT 2.0.4.

**Supplementary Table 14 | AI-assistant evaluation tasks and outcomes.** Success by agent and arm (A, skill and tools; B, command line and documentation; C, no tools), by task category, and release-guard re-test without and with the added input guards. Cross-judge agreement 0.913 (κ = 0.818).

**Supplementary Table 15 | Analyses with additive effects and no interaction: conditions and family-wise rejection rates.** Conditions and family-wise rejection rates (any group rejected) of the analyses with additive effects and no interaction.

## Supplementary Data

**Supplementary Data 1 | Calibration, power and comparator tables.** Sheets S2-1 to S2-11; the README sheet lists the content of each sheet.

